# The effect of flow on swimming bacteria controls the initial colonization of curved surfaces

**DOI:** 10.1101/866491

**Authors:** Eleonora Secchi, Alessandra Vitale, Gastón L. Miño, Vasily Kantsler, Leo Eberl, Roberto Rusconi, Roman Stocker

## Abstract

The colonization of solid surfaces by bacteria is a widespread phenomenon with major consequences on environmental processes, biotechnology and human health. While much is known about the molecular mechanisms of surface colonization, the influence of the physical environment remains poorly understood. Here we show that the magnitude and location of colonization of non-planar surfaces by motile bacteria is largely controlled by the local flow conditions. Using microfluidic experiments with *Pseudomonas aeruginosa* and *Escherichia coli*, we demonstrate that the velocity gradients created by a curved surface drive preferential attachment to specific regions of the collecting surface, namely the leeward side of cylinders and immediately downstream of apexes on corrugated surfaces, locations that are in stark contrast to where non-motile cells attach. The preferential attachment location depends on the local hydrodynamic conditions and, as revealed by a mathematical model benchmarked on the observations, on cell morphology and swimming traits, while it is independent of the physicochemical properties of the surface. The interplay between imposed flow and bacterial motility further affects the overall attachment rate, increasing it by up to two orders of magnitude compared to the non-motile case at moderate flow velocities of up to twenty times the bacterial swimming speed. These results highlight the importance of fluid flow on the magnitude and location of bacterial colonization of surfaces and provide a mechanistic model to predict colonization in flow for a wide range of applications, from infection biology to bioremediation and biofouling.

## Introduction

The attachment of individual bacteria to a surface is the first step toward the formation of surface-attached communities known as biofilms ^1^. Biofilms grow on human tissue, including the lungs, urinary tract, eyes, and in chronic wounds ^2^, and on implanted devices, such as catheters, prosthetic cardiac valves and intrauterine devices, posing serious health threats and reducing device lifetime ^3,4^. Biofouling and biocorrosion are ubiquitous, costly problems also in other settings, from industrial wastewater systems to marine environments ^5,6^. To date, the majority of research on bacterial colonization of surfaces has focused on flat surfaces ^7–12^; yet in many applications surfaces are not flat. As a result, no general framework exists to account for the role of surface shape on bacterial colonization.

Bacterial transport is often affected by fluid flow, a feature of many microbial habitats. In a straight microfluidic channel, motile bacteria become trapped close to flat surfaces, inducing a strong concentration of cells close to the channel walls ^12^. The trapping is a hydrodynamic phenomenon, determined by the action of fluid shear on motile, elongated bacteria ^12,13^. Bacterial accumulation also occurs in shallow microfluidic channels behind an obstacle and after a constriction^14,15^. In groundwater, the size of the grains of the porous matrix ^16,17^ and the heterogeneity in flow velocities ^17,18^ affect the transport of colloids and, potentially, bacteria. In the human body, secondary flows in the lungs depend on airway geometry ^19^, while urine transport is controlled by the amplitude of the contraction waves causing peristaltic motion in the ureter ^20^. In the gut, both the luminal flow and the undulated morphology of intestinal villi affect the adhesion to the gut epithelium and the growth of pathogenic bacteria, such as enteroinvasive *Escherichia coli*, which, in the absence of peristaltic motion, overgrow and trigger an immune response and inflammation ^21^. Hosts can also exploit flow-mediated transport of bacteria to favor bacterial adhesion to their surfaces, as occurs in the gut of the bobtail squid *Euprymna scolopes*, whose cilia create a flow that favors the recruitment of symbiotic *Vibrio fischeri* ^22^. Similarly, the ciliated epidermal surface of corals creates flows that stir the coral’s boundary layer, enhancing oxygen transport ^23^ and potentially affecting the transport of symbionts and pathogens. In these and many other scenarios, bacteria are recruited to surfaces that are not flat ^14,15,24–26^. Yet despite its biological relevance, the understanding of the influence of local flow conditions on the ability of bacteria to colonize uneven and curved surfaces has so far been limited.

The control of surface contamination by particulates in flowing fluids is important in many applications, such as filtration processes. The microscopic description of filtration usually considers a filter medium as an assembly of collectors, which capture particles suspended in the flow when these encounter one of the collectors ^27–30^. A frequent case, also relevant for many microbial processes, is the capture of micron-sized suspended particles by cylindrical collectors in laminar flow. In this case, the encounter can occur by either direct interception ^31,32^, when a particle travels on a streamline that passes sufficiently close (within one particle radius) to the collector to contact it, or diffusional deposition ^30,32^, when Brownian motion across the streamline causes the contact. When considering motile microbes rather than passive particles, motility can substantially enhance encounter rates ^33,34^, as shown for the colonization of marine organic aggregates ^35^. However, encounter rate studies to date have largely neglected the effect of flow on motility. Based on the strong effects of flow on motility observed in straight channels ^12^, we asked how flow shapes the encounter rate and encounter location of motile bacteria with curved surfaces.

In this work, we show that the attachment of motile bacteria to curved surfaces is controlled by the effect of the surface on the local flow. Using two model bacteria frequently found in environmental and clinical settings – *Pseudomonas aeruginosa* and *Escherichia coli* – we demonstrate that the flow conditions created by the curvature of a surface drive bacteria towards specific locations on the surface. We show that the interplay between local flow and bacterial motility affects both the attachment rate and the attachment site of bacteria, due to the deflection of the trajectories of swimming bacteria by the flow, and that this effect strongly depends on the magnitude of the flow. We present a mathematical model, which is in good agreement with the observations and provides a new tool to predict the location and magnitude of bacterial attachment to surfaces.

## Results

### Fluid shear affects the attachment rate of motile bacteria on a cylindrical pillar

As a prototypical case of a curved surface, we first consider the bacterial colonization of cylindrical pillars. This shape represents a simplified model system for grains in porous media ^36^, submerged benthic plants and filter fibers ^32,37,38^, and tissue heterogeneities in the body such as intravascular pillars ^39^. We found that the attachment rate of motile bacteria to a pillar strongly depends on the flow velocity and, in slower flow conditions, can be up to two orders of magnitude larger than for passive particles of the same size. This result was obtained by visualizing GFP-labeled *P. aeruginosa* PA14 swimming near and attaching to single pillars of different diameters (*d*_P_) exposed to different flow velocities (*U*) in a microfluidic device (Fig. 1a). Video microscopy was used in fluorescence mode to quantify bacterial attachment to the pillar (Fig. 1d) and in phase-contrast mode to capture bacterial trajectories.

**Figure 1.**
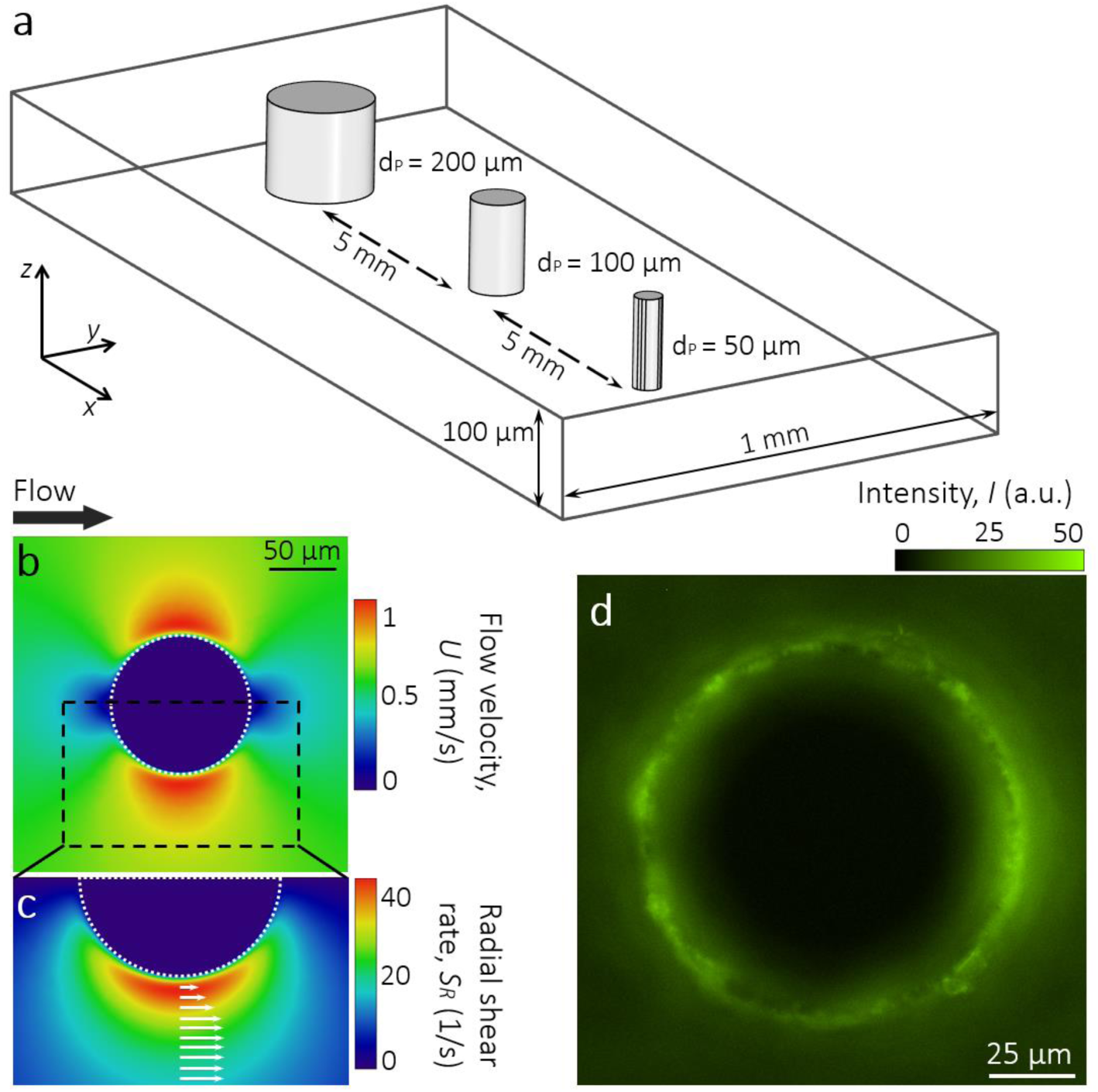
Microfluidic model of bacterial colonization on curved surfaces in the presence of fluid flow. **a** Schematic of the microchannel containing pillars of different diameters, *d*_P_ (200 μm, 100 μm and 50 μm; each repeated 2 times). **b-c** Flow velocity, *U* (b) and radial shear rate, *S*_*R*_ (c) around a 100-μm pillar, computed with COMSOL Multiphysics at a mean flow velocity of 500 μm/s. Superimposed arrows indicate the local velocity field. **d** Fluorescent image of GFP-tagged *P. aeruginosa* PA14 *wt* cells attached to a 100-μm pillar after flow at rescaled velocity *U/V* = 3.3 (*U* = 150 μm/s).

The role of motility was determined by comparing a motile strain (PA14 *wt*; Fig. 2a,f) with two non-motile mutants (PA14 *flgE*; PA14 *motB*; Fig. 2b,c,f). For the *d*_P_ = 100 μm pillar, the fluorescent intensity integrated over 5-μm-thick annulus around the perimeter of the pillar, *I*_IN_, is more than one order of magnitude higher for motile bacteria than for the two non-motile mutants at flow velocity *U* = 300 μm/s (Fig. 2f). This flow velocity is *U*/*V* = 6.6 times the mean bacterial swimming speed *V* (for PA14 *wt, V* = 45 ± 10 μm/s; Supplementary Fig. 1). In the following, we will use the rescaled flow velocity *U/V* to describe the relative magnitude of the imposed flow. This observation reveals that the strong increase in bacterial capture promoted by motility, already reported for sinking spheres ^33,34^, is valid also for pillars, when exposed to moderate flow (up to the threshold of *U/V* < 20, determined below). The mechanism of this enhanced capture is known: whereas passive particles and non-motile bacteria are captured only when the streamline along which they are transported comes within one particle or cell radius from the pillar ^28,29,32^, motile bacteria swim across streamlines and can reach the pillar surface from much larger distances ^33,34^ (Supplementary Fig. 2).

**Figure 2.**
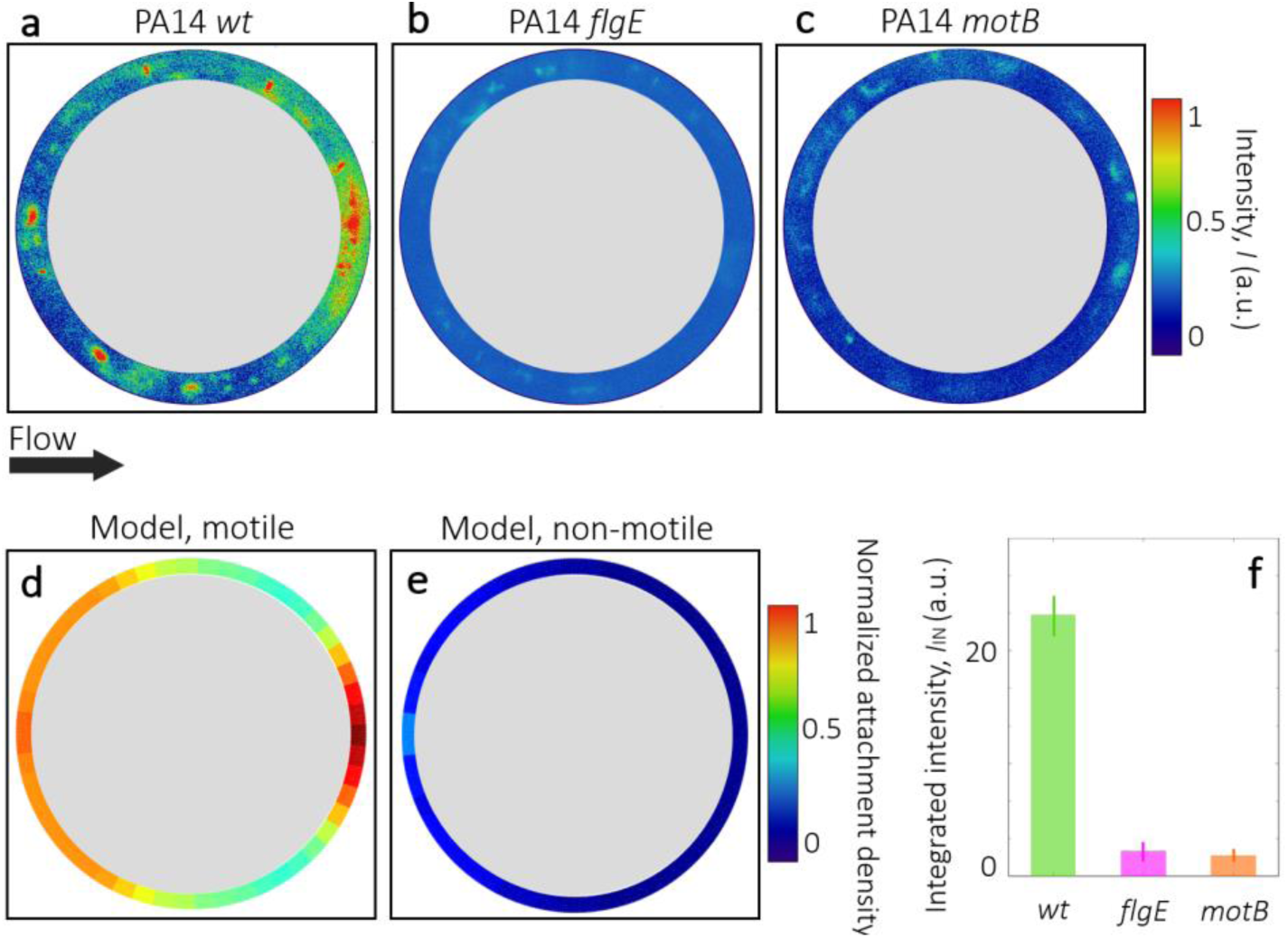
Surface colonization on pillars by *P. aeruginosa* is determined by bacterial motility. **a-c** Intensity distribution of the fluorescent signal from GFP-labelled motile (PA14 *wt*, a) and non-motile (PA14 *flgE*, b; PA14 *motB*, c) *P. aeruginosa* cells attached to a 100-μm pillar after flow at a rescaled flow velocity of *U/V* = 6.6. **d-e** Angular distribution of the normalized attachment density of bacteria on the pillar obtained with a mathematical model for motile (d) and non-motile (e) cells for the same flow rate and pillar dimension as a–c. **f** Integrated intensity, *I*_IN_ obtained for the motile (*wt*) and non-motile (*flgE, motB*) strains from the images in panels a–c.

The motility-induced increase in bacterial capture for moderate flow is correctly predicted by a mathematical model of bacteria swimming in flow (Fig. 2d,e). In the model, we first computed the three-dimensional velocity field in the microchannel using a finite element code (COMSOL Multiphysics), for the same geometry and flow rate as in the experiments (Fig. 1b,c; Methods). We then used this velocity field to compute the trajectories of single bacteria with an individual-based model (Fig. 4b, Supplementary Information; Methods). We modeled bacteria as prolate ellipsoids with aspect ratio *q* and swimming velocity *V* directed along the major axis. Their swimming direction at each instant in time was determined by a torque balance that accounts for the hydrodynamic shear from the flow and random fluctuations in the bacterial orientation due to rotational Brownian motion or tumbling, which are taken into account using an effective rotational diffusivity *D*_R_ ^12^. All parameters in the model (*q, V, D*_R_) were measured directly in separate experiments by tracking individual cells in the absence of flow (Methods; Supplementary Fig. 1). The model scored all contacts with the surface of the pillar (neglecting steric or hydrodynamic interactions) to obtain a capture rate. The capture efficiency, *η*_C_^mod^, was computed as the fraction of bacteria removed from the volume of water passing through the region defined by the pillar’s cross section, following the classic definition ^28,32^. The capture efficiency is equivalent to the capture rate divided by the flux of bacteria passing through the cylinder ^32^ and can be directly compared to its experimental counterpart, *η*_C_^exp^, which was obtained from the integrated fluorescent intensity *I*_IN_ (Methods).

**Figure 3.**
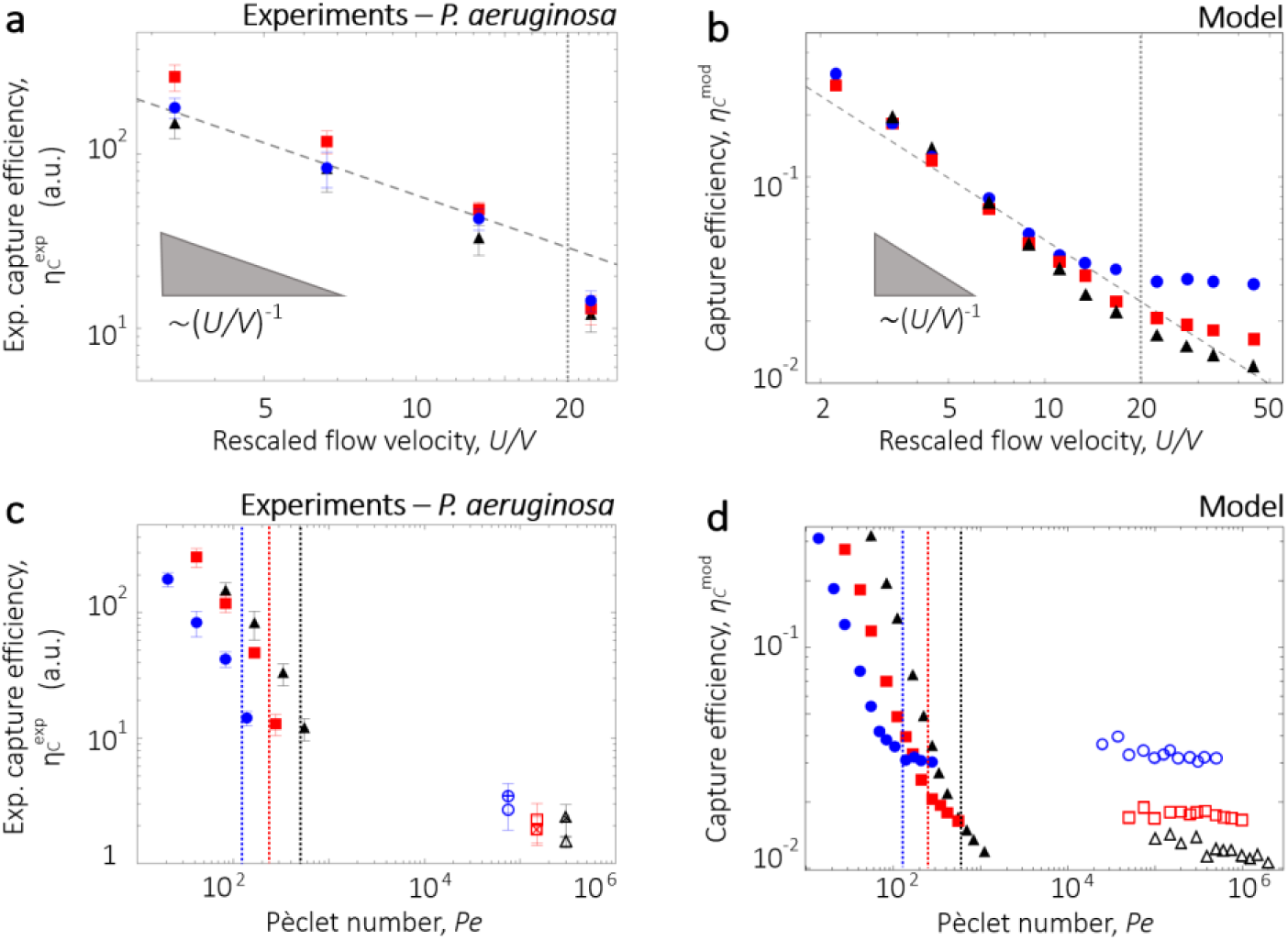
Capture efficiency of *P. aeruginosa* depends on pillar dimension and flow velocity. **a** Experimental capture efficiency, *η*_C_^exp^, of motile P. aeruginosa PA14 *wt* cells, as a function of the rescaled flow velocity *U/V*, for pillars of diameter 50 μm (blue circles), 100 μm (red squares) and 200 μm (black triangles). The dashed curve shows the scaling 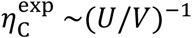. **b** Capture efficiency, *η*_C_^mod^, as a function of *U/V*, obtained from the model for the same pillar diameters as in a. **c** Experimental capture efficiency, *η*_C_^exp^, as a function of the Péclet number, *Pe*, for motile PA14 *wt* (filled symbols) and non-motile (PA14 *flgE*, open symbols with a cross; PA14 *motB*, open symbols) cells, for pillars of different diameters. **d** Capture efficiency, *η*_C_^mod^ as a function of the *Pe*, obtained from the model for different pillar diameters, in the case of motile (filled symbols) and non-motile (open symbols) cells. Vertical dotted lines in panel c and d represent the *Pe* corresponding to *U/V* = 20, calculated for each pillar dimension.

**Figure 4.**
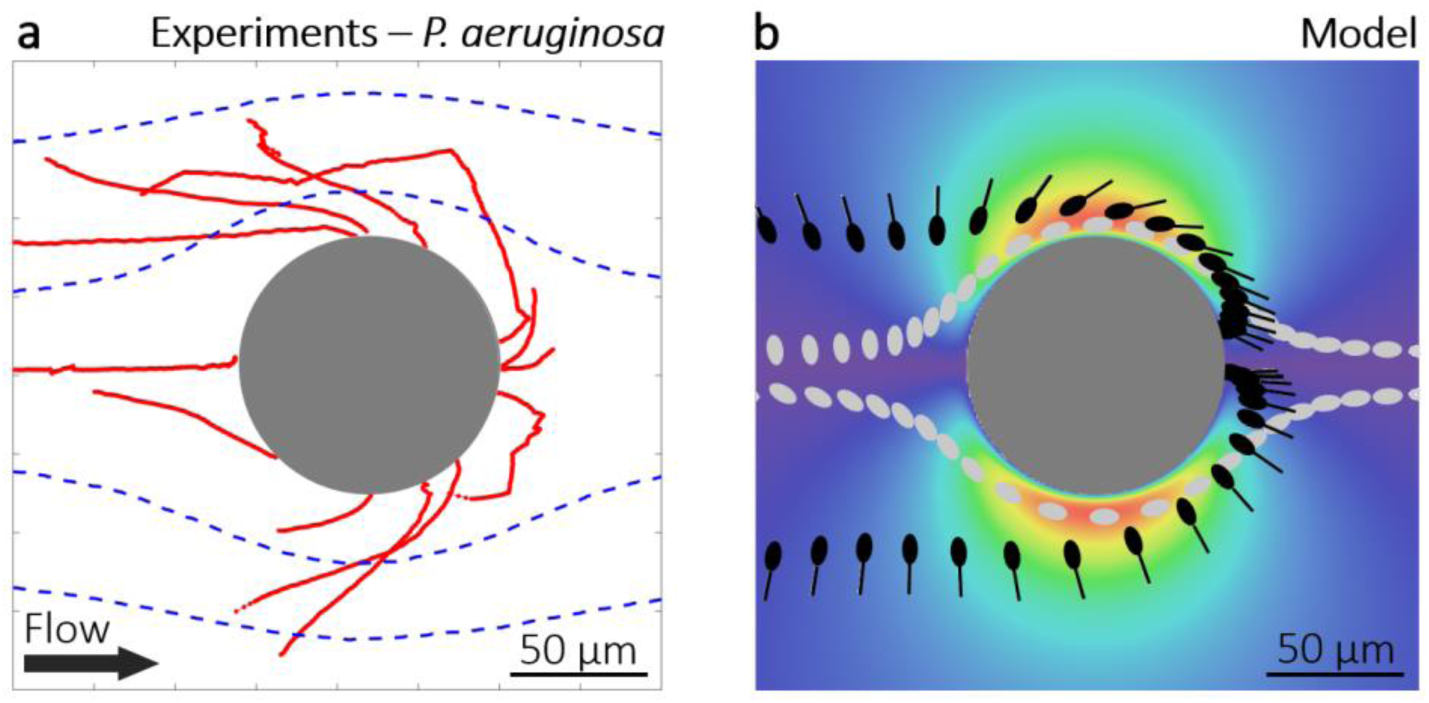
Fluid shear affects the trajectories of swimming bacteria around a pillar. **a** Sample trajectories of *P. aeruginosa* PA14 *wt* cells in flow around a 100-μm pillar at *U/V* = 3.3 (*U* = 150 μm/s), for motile (red lines) and non-motile (blue dashed lines) cells. **b** Trajectories of motile (black) and non-motile (gray) cells, simulated without rotational noise ξ_R_, in flow around a 100-μm pillar at *U/V* = 3.3 simulated with the model. The colour scale represents the radial shear rate, *S*_*R*_, around a 100-μm pillar at *U* = 500 μm/s, reported in Fig. 1c.

Two regimes emerge when considering the capture efficiency of motile cells as a function of the imposed flow – one regime for moderate flows (*U/V* < 20) and a different regime for strong flows (*U/V* > 20). In the moderate flow regime, the capture efficiency is found to be strongly dependent on fluid velocity, but independent of the pillar diameter: the capture efficiencies *η*_C_^mod^ and *η*_C_^exp^ as a function of *U/V* collapse onto a single curve for different pillar diameters (Fig. 3a,b). This means that for moderate flows, the attachment rate and therefore the density of attached bacteria are the same regardless of pillar diameter (Fig. 3a,b). In this regime, we observe a scaling dependence of both *η*_C_^mod^ and *η*_C_^exp^ with (*U*/*V*)^−1^ (Fig. 3a,b), indicating that the capture efficiency depends inversely on the fluid velocity (Supplementary Fig. 3a) and linearly on the bacterial swimming speed (Supplementary Fig. 3b). In contrast, in the strong flow regime (*U/V* >20) the predicted capture efficiency *η*_C_^mod^ does not depend on fluid velocity (Fig. 3b) and decreases with increasing pillar diameter (Fig. 3b). As a result, the attachment density of attached bacteria is higher for smaller pillars, as for the case of passive particles ^32^.

The strong increase in the attachment rate of motile compared to non-motile bacteria can be further seen by considering the capture efficiencies *η*_C_^mod^ and *η*_C_^exp^ as a function of the Péclet number, *Pe = Ud*_P_*/D*, which provides a measure of the importance of transport by flow relative to transport by diffusion. Here *D* is the translational diffusion coefficient of the bacteria, given by the Brownian diffusivity of the cells for non-motile bacteria or by the effective diffusivity due to motility for motile bacteria, with the latter approximately three orders of magnitude higher than the former ^40^ (Supplementary Information). In our experiments and simulations, motile bacteria are thus characterized by lower values of *Pe* than non-motile bacteria, given their higher values of *D*. In the low-*Pe* regime, the enhancement in the capture efficiency is apparent, and can be understood as diffusive transport being important relative to transport by flow, i.e., motile bacteria being able to cross the fluid streamlines owing to their large diffusivity. For increasing flow rate, *Pe* increases and the capture efficiency of motile bacteria rapidly decreases as the role of transport by flow increases over transport by diffusion, until *Pe* is so large (corresponding to the strong flow regime, *U/V* > 20) that diffusive transport is overcome by transport due to fluid flow. (Fig. 3c-d, vertical dotted lines). In the latter regime, attachment rates become comparable with the values for non-motile bacteria and the capture efficiencies are nearly independent of *Pe* (Fig. 3d). We note that cell shape plays only a modest role in determining the overall capture rate in the regime we investigated (Supplementary Fig. 5l), implying that the overall capture rate is controlled primarily by the swimming speed and not by the torque exerted by the local shear rate.

### Fluid shear promotes the leeward attachment of bacteria to cylindrical pillars

Motility in flow affects not only the magnitude of attachment but, importantly, also the location of attachment. We demonstrate this first by considering again the case of a cylindrical pillar, both through tracking of individual bacteria before they contact the pillar, and by quantifying the spatial distribution of attachment on the pillar. At flow velocities that are up to a few times the bacterial swimming speed (*U/V* ≈ 3–6), tracking of individual *P. aeruginosa* cells in flow revealed trajectories directed towards the leeward side of the pillar (Fig. 4a, red line). These trajectories can be explained in terms of the effect of fluid flow on swimming cells. The no-slip condition on the surface of the pillar creates local velocity gradients (here for brevity termed ‘shear’) (Fig. 1b,c). Shear induces a hydrodynamic torque on bacteria, tending to rotate them ^41^. When bacteria are non-motile, this rotation is rather inconsequential, as they simply follow the flow streamlines (Fig. 4a, blue dotted paths). Instead, when they are motile, it redirects their trajectory ^12^ and causes them to reach the leeward side of the pillar (Fig. 4a, red paths). Because bacteria are preferentially aligned with streamlines (pointing either upstream or downstream) as they are transported past the pillar, the local shear created by the pillar induces a torque that directs bacteria pointing downstream towards the leeward side of the pillar (Fig. 4a, red paths) and bacteria pointing upstream away from the pillar. In contrast, non-motile cells simply follow the flow streamlines (Fig. 4a, blue dotted paths).

This mechanism of inward-focusing at the hands of the local shear created by the curved surface is confirmed by our mathematical model. The model further shows that, for conditions mimicking those in the experiments, this inward-focusing is sufficiently rapid to cause bacteria to contact the pillar on its leeward side. An important element here is the low-flow velocity in the wake of the cylinder: once bacteria are redirected into that region, their swimming speed will often be larger than the local flow velocity, making upstream swimming possible and enabling attachment of cells to the leeward side of the pillar (Fig. 4b, black cells). We note that this process does not consider sensing or directed motion toward the surface, which were not included in the model: leeward contact is driven purely by the deflection of swimmers’ trajectories at the hands of the local shear. Finally, even non-motile bacteria are reoriented by shear, but this reorientation does not result in net movement towards the pillar (Fig. 4b, gray cells).

By redirecting cells towards the leeward side of the pillar and consequently increasing the number of local contact events with the surface, the interaction between shear and motility promotes the preferential colonization of specific regions of the pillar’s surface. The heterogeneity in surface colonization is confirmed by our observations of cell accumulation around the perimeter of the pillar, measured in terms of the fluorescence intensity of motile, GFP-tagged PA14 *wt* cells (Fig. 1d; Supplementary Information). The preferential leeward attachment is clearly visible in terms of a higher fluorescence intensity, for example for the case of a flow velocity *U/V* = 6.6 (Fig. 1d, 2a). Preferential leeward attachment was observed only for motile cells, and was not observed for the two non-motile PA14 strains (*flgE* and *motB;* Fig. 2b, c). The latter preferentially attach on the windward side of the pillar, as expected for passive particles ^32^. In agreement with the experimental evidence, the mathematical model shows that bacterial motility determines the distribution of bacterial attachment sites around the pillar: motile cells preferentially attach on the leeward surface of the pillar (Fig. 2d), whereas non-motile cells attach only on its windward side (Fig. 2e). Furthermore, in experiments in which the flow direction was reversed after 2.5 h, a second, symmetric attachment peak formed on the opposite side (the originally windward side) of the pillar over the ensuing 2.5 h (Supplementary Fig. 4), independently of the growth medium (Supplementary Fig. 4), further confirming the robustness of the phenomenon.

The flow velocity affects the distribution of attachment of motile bacteria around the pillar, with higher flow velocities shifting the attachment from the leeward to the windward side. Leeward attachment was observed for motile cells in moderate flows (*U*/*V* < 20; Fig. 2a). In contrast, for strong flows (*U*/*V* > 20), motile bacteria attached preferentially to the windward side (Fig. 5), as passive particles do ^32^. The shift in the location of preferential attachment as a function of the flow velocity can be quantified by computing the angular distribution of the attachment sites on the surface of the pillar. These distributions were obtained from the experiments by using the fluorescence intensity signal of the GFP-labelled bacteria attached along the perimeter of the pillar (blue lines in Fig. 5a-d) and from the model by using the distribution of contact sites obtained from the trajectories of 10^5^ bacteria (orange lines in Fig. 5a-d). The angular distributions of bacterial attachment accord in the model and experiments and show a clear dependence on flow velocity. At a low flow velocity of *U/V* = 3.3 we observed an accumulation region on the leeward side of the pillar (Fig. 5a), which becomes more pronounced when the flow velocity is doubled (*U/V* = 6.6; Fig. 5b, f). A further increase in the flow velocity (*U/V* = 13.3) shifts the preferential attachment toward the windward side of the pillar and leaves just a small leeward peak (Fig. 5c), which completely disappears at *U/V* = 22.2 (Fig. 5d,g). This shift is consistent with the observed transition from a motility-dominated to a flow-dominated regime with increasing flow velocity (Fig. 3).

**Figure 5.**
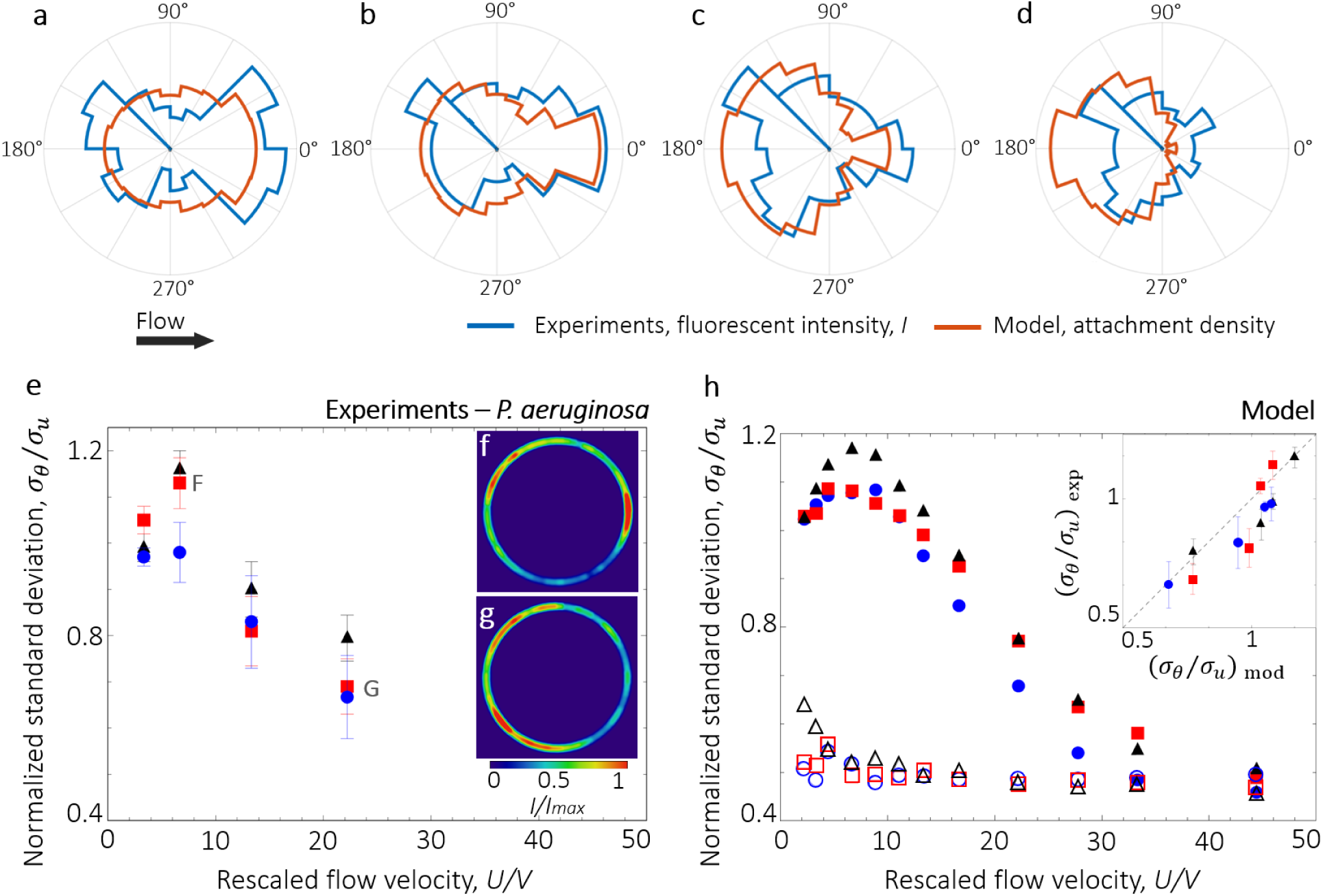
Fluid velocity modifies the angular distribution of bacterial colonization around a pillar. **a-d** Angular distribution of the fluorescence intensity, *I* (blue; experiments after 5 h of flow of a diluted suspension of PA14 *wt* GFP cells) and the simulated attachment density (orange; model) on a 100-μm pillar for a mean rescaled flow velocity *U/V* of 3.3 (a), 6.6 (b), 13.3 (c) and 22.2 (d). **e** Normalized standard deviation *σ*_*θ*_/*σ*_*u*_ for motile PA14 *wt* bacteria as a function of the rescaled flow velocity, *U/V*, for pillars of diameter 50 μm (blue circles), 100 μm (red squares) and 200 μm (black triangles). **f-g** Intensity distribution *I* of PA14 *wt* cells, normalized by the maximum value (*I*_max_), attached to a 100-μm pillar at *U/V* = 6.6 (f) and *U/V* = 22.2 (g). At moderate flows, preferential colonization occurs on the leeward side of the pillar. Flow direction is from left to right. **h** Normalized standard deviation *σ*_*θ*_/*σ*_*u*_ obtained with the model as a function of *U/V* and for the same pillar dimensions as in e. Results for motile cells (filled symbols) and for non-motile cells (open symbols) are shown. (Inset) *σ*_θ_/*σ*_u_ of the experimental intensity angular distribution as a function of *σ*_θ_/*σ*_u_ from the model for different pillar diameters. The dashed line represents *y = x*.

The change in the distribution of attachment of motile bacteria with flow velocity can be quantified in terms of the standard deviation, *σ*_*θ*_, of the angular distribution, normalized by the standard deviation, 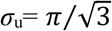, of a uniform angular distribution ^38^. Values of *σ*_θ_*/σ*_u_ > 1 denote a distribution skewed towards the leeward side of the pillar, whereas values of *σ*_*θ*_*/σ*_*u*_ < 1 denote preferential colonization of the windward side. This analysis reveals that the skewness of attachment in the leeward direction is most pronounced for *U/V* = 6.6 (*σ*_θ_*/σ*_u_ > 1) and diminishes at *U/V* = 22.2 (*σ*_θ_*/σ*_u_ = 0.6) (Fig. 5e,f), as the role of bacterial motility decreases and cells are primarily transported by the flow (Fig. 5e,g), akin to passive particles (*σ*_θ_*/σ*_u_ = 0.37) ^38^. The fact that the passive-particle limit is not reached in the experiments even for the highest flow velocity tested (1000 μm/s) implies that even when the flow speed is substantially higher than the swimming speed (*U*/*V* = 22.2), the process of attachment is not completely driven by flow, but still influenced by motility. In addition, the model for motile cells (Fig. 5h, filled symbols) shows the transition from leeward to windward attachment as the flow velocity increases from 2.2*V* to 22.2*V*, in very good agreement with observations (Fig. 5i), and also reaches the passive direct interception limit (*σ*_θ_*/σ*_u_ = 0.37), at a flow velocity *U/V* = 33.3. In the model, no dependence of *σ*_θ_*/σ*_u_ on the flow velocity is observed for non-motile bacteria (Fig. 5h, open symbols), confirming that the transition from leeward to windward attachment is determined by the interplay between bacterial motility and fluid flow. This transition in the bacterial distribution corresponds to the motility-dominated to flow-dominated transition observed in the capture efficiency for *U/V* > 20 (Fig. 3b): the interaction of motility and flow thus causes major differences in the capture of motile and non-motile bacteria not only in terms of the magnitude of bacterial attachment to surfaces, but also in terms of the spatial distribution of attachment.

Cell shape is a key determinant of the inward-focusing and leeward encounter process. While self-propulsion is necessary for a cell to reach the pillar surface once in the leeward stagnation region, it is the elongated shape of bacteria that is responsible for their preferential alignment with the flow direction during transport past the cylinder, and hence their inward motion toward the leeward stagnation region. This preferential alignment is well known in the context of Jeffery orbits ^12,42^, i.e., the periodic rotation of spheroids in laminar flow: the more elongated the spheroid, the larger the fraction of time it spends aligned with streamlines. The role of elongation is confirmed by results from our model, showing that swimming spherical cells, which reach the leeward stagnation region with a random orientation, have a lower attachment density to the leeward surface of the pillar (Supplementary Fig. 5). We therefore expect that spherical swimmers, such as the alga *Chlamydomonas reinhardtii* ^43^, will not exhibit preferential leeward encounters through this mechanism (Supplementary Fig. 5e-i). The vast majority of motile bacteria, however, are elongated, either because of the elongation of their cell body, or even more so because of the presence of flagella that enhance their effective aspect ratio ^12^. Numerical results predict a sharp increase in the leeward attachment when swimmer elongation increases from 1 (spherical) to 3, and a plateau in the attachment for further increases in elongation (Supplementary Fig. 6).

A further trait affecting the transport of bacteria towards the pillar is the frequency of the cells’ own reorientation (or ‘tumbling’), which can be expressed in terms of a rotational diffusivity, *D*_*R*_^12^. Simulations show that cells with low rotational diffusivity (*D*_*R*_ = 10^−2^ rad^2^ s^−1^; ‘smooth swimmers’) have a stronger tendency for leeward attachment compared to cells with high rotational diffusivity (*D*_*R*_ = 1.4 rad^2^ s^−1^; value obtained for *P. aeruginosa*, ‘tumbling cells’, Supplementary Fig.1) (Supplementary Fig. 6). A further (though unrealistically high) increase in rotational diffusivity (*D*_*R*_ = 10 rad^2^ s^−1^) results in preferential windward attachment. These results support the hypothesis that alignment along streamlines and reorientation by local shear drive leeward attachment, whereas random fluctuations in the orientation of bacteria such as those caused by tumbling diminish it. Moreover, even for smooth swimmers the model predicts the transition from leeward to windward attachment with increasing flow velocity, but with colonization spots peaks that are more pronounced compared to the ones obtained for tumbling cells (Supplementary Fig. 7).

Leeward attachment originates from the interaction between motility and the shear generated by the pillar, not from the physico-chemical properties of the pillar. We fabricated PDMS pillars of different wettability (Fig. 6a) and stiffness (Fig. 6d), and tested these for a single flow rate (*U*/*V* = 6.6; Fig. 6). Whereas surface coverage varied with surface properties (being lower for more hydrophilic and softer surfaces; Fig. 6c, f), the angular distribution of surface colonization was similar and a leeward attachment peak was robustly observed for all pillars tested (Fig. 6b, e). Similarly, leeward attachment is not a consequence of the use of a specific medium. Growth media can affect bacterial adherence to surfaces, due to changes in pH and electrolyte concentration^44^. We quantified the attachment distribution of bacteria in two media (TB and AB minimal medium) and for different dilutions of the same medium (Supplementary Information). Results show that the bacterial distribution was similar regardless of the medium used and the leeward attachment was always observed (Supplementary Fig. 7). The use of a diluted medium allows us to exclude that bacterial growth was responsible for the observed angular distribution of bacteria around the pillar, as growth in the diluted medium would have been too slow. Taken together, these observations confirm the purely hydrodynamic nature of the leeward attachment phenomenon, which does not depend on the physico-chemical properties of the surface or on the adhesiveness of bacteria. The increase in bacterial density in preferential areas of the pillar is due to a flow-induced increase in the probability of contact between the bacterium and the surface, thus creating a preferential colonization spot.

**Figure 6.**
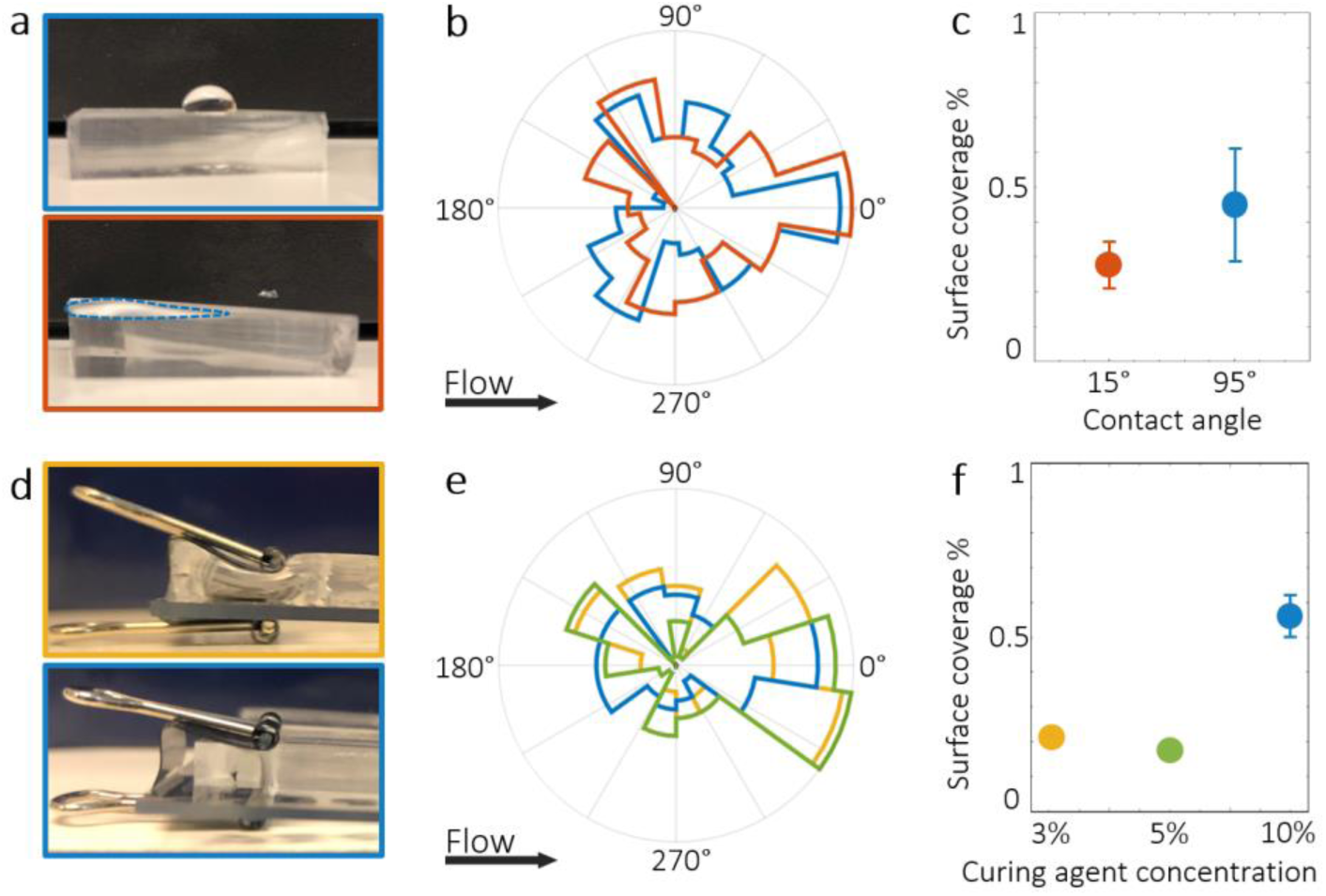
Surface properties of the pillar do not affect the angular distribution of bacterial colonization. **a** Tryptone broth droplets on PDMS surfaces 5 days (upper panel; contact angle 95° ± 5°) and 1 hour (lower panel; contact angle 15°± 5°) after plasma treatment showing, respectively, the hydrophobic and hydrophilic nature of the two surfaces. In the lower panel, the droplet wets the surface creating a film: its borders are marked with a blue dashed line in the image for ease of visualization. **b** Angular distribution of the fluorescence intensity, *I*, of PA14 wt GFP cells attached on a hydrophobic pillar (blue) and on a hydrophilic pillar (orange), for a flow velocity at *U/V* = 6.6 and a pillar diameter of 100 μm. **c** Surface coverage (measured on the upper surface of the microfluidic channel) for the hydrophobic surface (blue) and the hydrophilic surface (orange) under the same experimental conditions as panel b. **d** Slabs of PDMS containing 3% curing agent (upper panel; Young modulus = 150 ± 50 kPa) and 10% curing agent (lower panel; Young modulus = 2.25 ± 0.25 MPa), respectively, undergoing compression from a paper clip in order to visualize the difference in stiffness. **e** Angular distribution of the fluorescence intensity, I, of PA14 wt GFP cells attaching on a pillar containing 3% (yellow), 5% (green) and 10% (blue) curing agent, for a flow velocity at *U/V* = 6.6 and a pillar diameter of 100 μm. **f** Surface coverage (measured on the upper surface of the microfluidic channel) for PDMS containing different concentrations of curing agent under the same experimental conditions as panel e.

### Fluid shear promotes the leeward attachment of bacteria to general corrugated surfaces

The flow-induced, preferential leeward attachment of motile bacteria to curved surfaces is a more general phenomenon, characterizing attachment on uneven surfaces more generally and for different bacterial species. We demonstrate this by showing the spatial attachment pattern of YFP-labelled *E. coli* (strain HCB1733) in a microfluidic channel with sinusoidally-shaped sidewalls (50 μm wavelength, 25 μm amplitude; Fig. 7a), and extending our mathematical model to this geometrical configuration. Bacterial surface colonization was again quantified in terms of fluorescence intensity in the experiments and contact events with the surface in the model. At moderate flows (*U* = 150 μm/s; *U/V* = 6.9, calculated using the swimming speed of *E. coli, V* = 21.6 μm/s; ^25^), bacteria attached to the channel sidewalls preferentially in the region immediately following the apexes of the sinusoid, both in the experiments (Fig. 7b) and the model (Fig. 7c). Preferential attachment leeward of apexes is shown, for example, by the peak in the fluorescent intensity near the location *x* = 20 μm of a period of the sinusoidal surface, obtained experimentally by averaging over 100 identical periods (Fig. 7b) and confirmed by modeling results (Fig. 7c). The mechanism is the same as for pillars: elements of a curved surface that protrude into the flow – in this case, the apexes of the sinusoid – generate a locally higher shear (Fig. 7d), which rotates bacteria and drives motile bacteria toward the surface. As a result, they preferentially contact the surface downstream of the protruding element, if their swimming speed is greater than the local flow velocity (Fig. 7e, yellow cell). These results thus suggest that our findings can be generalized: fluid shear promotes the preferential attachment of motile bacteria to the leeward side of curved elements of a surface.

**Figure 7.**
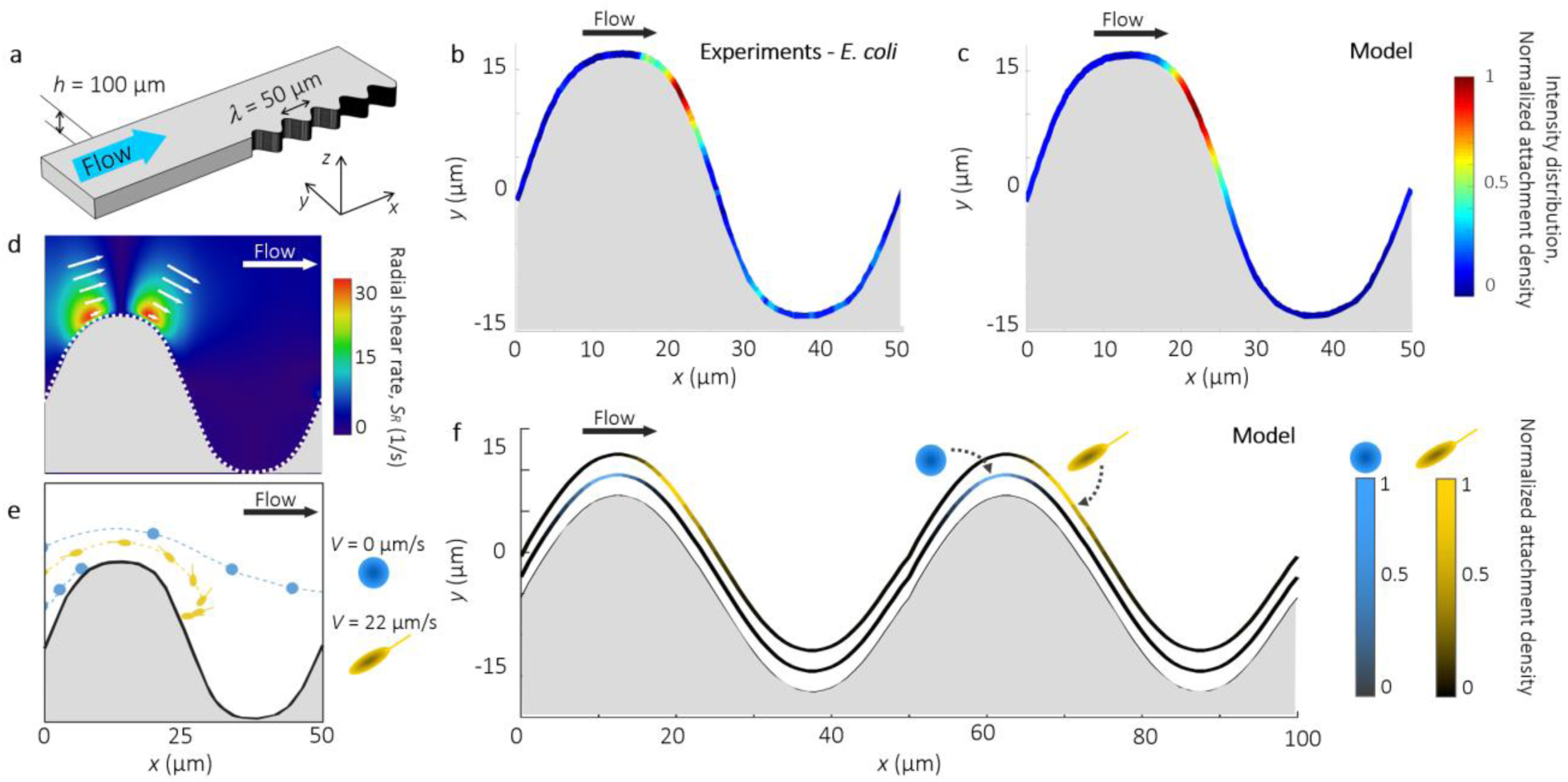
Preferential leeward attachment of a sinusoidal surface by motile *E. coli*. **a** Schematic of the microchannel with a sinusoidal lateral surface with wavelength 50 μm and amplitude of 25 μm. **b** Intensity distribution of the fluorescence intensity from GFP-tagged *E. coli wt* cells attached to the sinusoidal surface after 3 h of flow at *U* = 150 μm/s (*U/V* = 6.9), averaged over 100 identical periods and normalized for the maximum intensity value. **c** Normalized attachment density of cells on the sinusoidal surface obtained from the model at *U* = 150 μm/s. **d** Radial shear rate, *S*_*R*_, around one period of the sinusoidal lateral wall, computed with a finite element method at *U* = 500 μm/s. Superimposed arrows indicate the local velocity field. **e** Simulated trajectories of spherical non-motile (blue) and of elongated motile (yellow) cells in flow around a period of the sinusoidal wall. **f** Simulated attachment density of spherical non-motile and motile bacteria on the sinusoidal wall at *U* = 150 μm/s.

Surface regularity is not a prerequisite of leeward attachment. To illustrate this, we carried out microfluidic experiments with an irregularly corrugated surface (Fig. 8), a model for surface roughness in natural and artificial microbial environments. Experimental observations and modeling results both again show strong heterogeneity in surface colonization by motile bacteria (Fig. 8). The model also shows that this heterogeneity is absent for non-motile bacteria (Supplementary Fig. 9). Preferential attachment after apexes can be even more pronounced than for the sinusoidal surface, as indicated by both the experimental data (Fig. 8a,b) and the model (Fig. 8c). Examples include the peaks in the fluorescent intensity near the locations *x* = 450, and 700 μm (Fig. 8). This is in line with the fact that an irregular surface is composed of elements with significantly different curvature, each of which affects the flow (and thus the shear) differently. Despite the topographical complexity of this surface, the model also predicts strong preferential attachment locations, with reasonable agreement with the observations. The complexity of this configuration, which we suggest partly accounts for the differences between observations and model, stems from the fact that the effect of different surface elements on bacterial trajectories is not independent, but rather a bacterial trajectory integrates all the effects of successive curvature elements. Fundamentally, however, observations and the model support both the existence and the leeward localization of preferential attachment on generally shaped surfaces.

**Figure 8.**
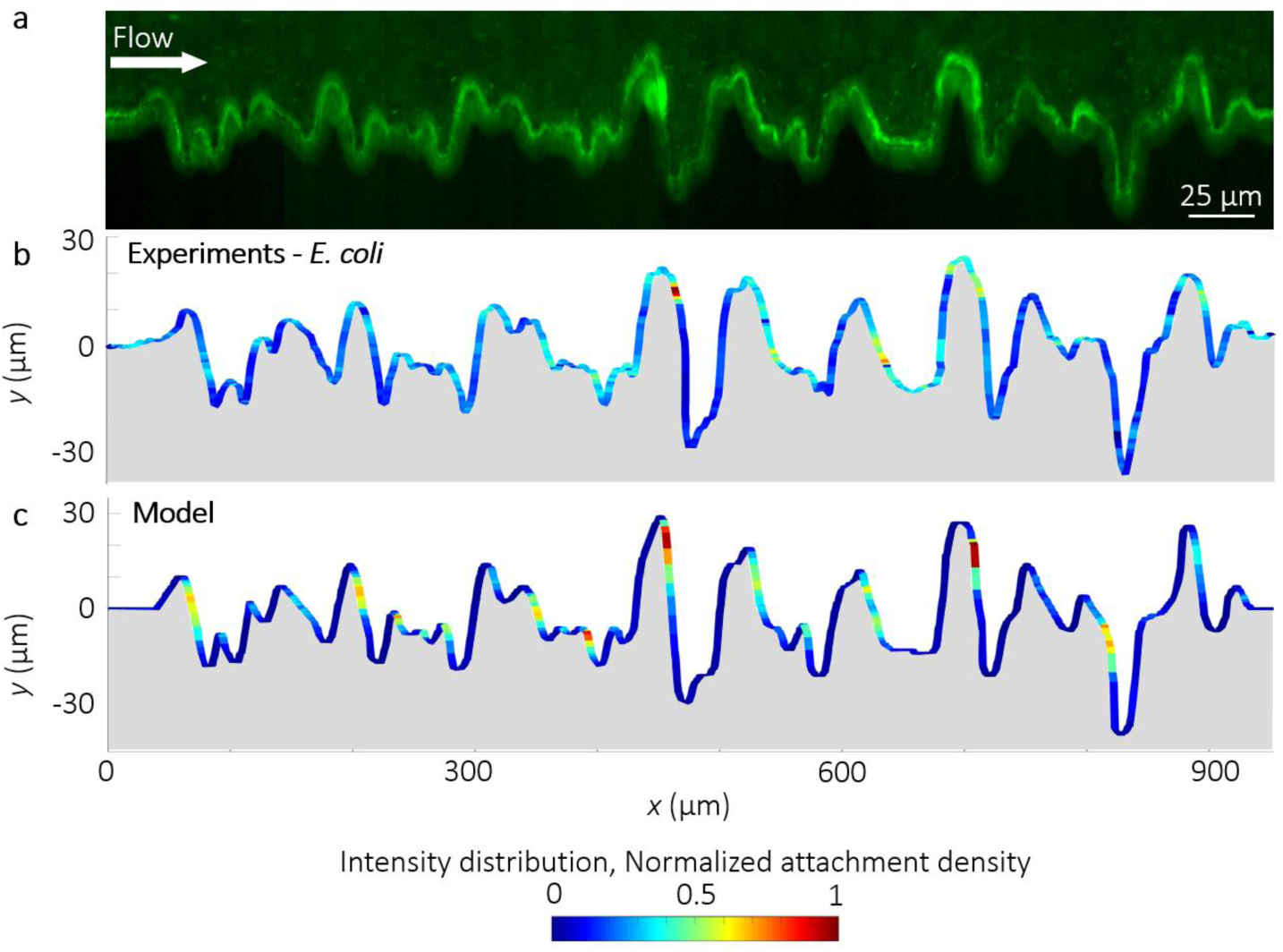
Colonization of a randomly corrugated surface by *E. coli*. **a** Fluorescent image acquired at channel mid-plane of GFP-tagged *E. coli wt* cells attached to the lateral corrugated surface of a microfluidic channel (height = 100 μm) after 3 h of flow of a diluted bacterial suspension at a mean flow velocity of *U* = 150 μm/s. **b** Intensity distribution of the fluorescent signal shown in panel a, normalized by its maximum value. **c** Normalized attachment density of cells (elongation *q* = 8.5, swimming speed *V* = 21.6 μm/s) on the corrugated surface obtained with the model at a mean flow velocity of *U* = 150 μm/s (*U/V* = 6.9).

## Discussion

The role of bacterial motility in favoring contact with and thus attachment to surfaces has long been known ^45^, and is considered an important early step in the transition from a planktonic state to the surface-associated state that initiates biofilm development. However, the role of fluid flow on this process has rarely been considered from a mechanistic viewpoint, with regard to determining how the effect of flow on the trajectories of bacteria influences surface colonization. For flat surfaces, flow has been proposed to favor surface colonization through shear trapping ^12^. Here we focused on curved surfaces and showed two major consequences of flow. First, in the presence of flow, bacterial motility strongly enhances the attachment of bacteria to surfaces. Second, fluid flow – in concert with bacterial elongation and swimming traits (speed, tumbling rate) – determines where a cell will contact a curved surface, resulting in surface colonization patterns and in particular leeward attachment that are strikingly different from those expected when ignoring the effect of flow or motility. The former observation extends results, mostly theoretical, obtained for the colonization of spherical aggregates, primarily in the context of sinking particles in the ocean ^33,34^, to cylindrical pillars of different dimensions and to a wide range of flow rates, which has particular relevance for applications ranging from filtration processes to biofouling and bioclogging. The second observation, to the best of our knowledge, has not been reported before, yet our experiments with different surface geometries, adherence properties and bacterial species indicate that the phenomenon of leeward attachment is likely to be very general.

The magnitude of fluid flow plays a fundamental role on leeward attachment. Once bacteria have been reoriented by flow, leeward attachment occurs if bacteria can swim against the flow in the wake of the pillar or other curved surface. When flow is too high, this cannot occur. Therefore, leeward attachment is not observed at flow velocities that are much higher than bacterial swimming speed. Here we observed leeward attachment for flow velocities as high as 13-fold the bacterial swimming speed, or 600 μm/s (for *P. aeruginosa*, which swim at 45 μm/s). While the exact threshold value of flow velocity below which leeward attachment is expected depends on the bacterial swimming velocity and elongation, flow velocities of a magnitude at which leeward capture is predicted occur very frequently in natural bacterial habitats. For example, in the soil, flow velocities for groundwater can vary between 1 and 1000 μm/s, depending on the soil type ^46,47^.

The process of shear-induced reorientation that leads to leeward attachment depends on two bacterial phenotypes: shape and motility. The role of these two phenotypes was already recognized in the related process of shear-induced trapping ^12^ and was here confirmed by our mathematical model for leeward attachment. A direct consequence of this dependence is a potential niche differentiation of bacterial species having different phenotypic traits: for a given surface geometry and flow velocity, bacteria with different shapes or different swimming characteristics will attach to different regions of the surface. To illustrate this, we used the model to predict the colonization of a sinusoidally corrugated surface by two different bacteria: a spherical, non-motile bacterium (Fig. 7e, blue) and an elongated, motile bacterium (Fig. 7e, yellow). The results show that flow causes non-motile bacteria to attach preferentially windward of apexes and motile bacteria preferentially leeward of apexes on the sinusoidal corrugation (Fig. 7f).

Since attachment sites are the seeding ground for biofilm formation, the interaction of flow and motility described here provides a mechanism for heterogeneous seeding of surfaces by bacteria with different phenotypic traits. Heterogeneity therefore can arise in a biofilm not only during development, due to spatial segregation driven by biological interactions, but can be present *ab initio* due to the physics of how bacteria encounter surfaces. Cell morphology has already been shown to be beneficial for surface colonization in flow in *Caulobacter crescentus*, suggesting that specific shapes can favor bacteria in different hydrodynamic conditions ^11^. At the community level, this interplay between phenotypic traits and flow could determine long-term population dynamics, akin to the flow-induced spatial segregation of *Vibrio cholerae* strains based on their different adhesiveness on flat surfaces ^48^. Our results suggest that in environments characterized by flow, niches of bacterial colonization of surfaces may be a function of cell morphology and swimming behavior, and that the flow environment can significantly affect bacterial meta-population dynamics by creating a feedback between flow conditions, surface topography and competition among species ^48,49^. This preferential attachment can thus play an important role in determining the structure, adaptation and potentially the evolution of microbial communities in aqueous ecosystems ^50^, and in medical ^21,22^ and industrial settings ^5,6^.

We have shown that flow–motility interaction can favor the formation of colonization hotspots on curved surfaces. Colonization hotspots, in turn, are favorable sites for biofilm formation and quorum sensing ^51–54^. Strong flow can repress quorum sensing, by diluting the concentration of inducer molecules, but the effect of flow is strongly quenched in sheltered regions, such as nooks and crevices ^51^. Leeward attachment hotspots may promote quorum sensing both due to the higher local bacterial density and by the sheltered nature of the attachment location. Since antibiotic resistance and pathogenicity are behaviors mediated by quorum sensing ^53,55^, this result highlights the potential importance of flow, motility and surface geometry in a wide range of health-related processes, with implications for the design of filters and medical devices. Moreover, the observation that bacterial transport and attachment is largely controlled by bacterial morphology, surface topography and flow demonstrates a previously unidentified and potentially ubiquitous interaction contributing to surface colonization in fluid environments. This knowledge, and the quantitative mechanistic model we have proposed, opens new frontiers in the possibility of controlling the colonization of surfaces by bacteria, and calls for a better understanding of the ecological and technological consequences in the many applications where the formation of biofilms is either desirable, such as in wastewater treatment plants and bioremediation systems, or to be avoided, such as artificial implants, medical devices and desalination membranes.

## Methods

### Bacterial cultures

Experiments were performed using GFP-tagged *P. aeruginosa* strain PA14 wild type, flagella-deficient strain PA14 *flgE* and motility-deficient strain PA14 *motB*, and GFP-tagged *E. coli* strain MG1655 wild type. *P. aeruginosa* and *E. coli* solutions were prepared by inoculating 3 mL Tryptone Broth (TB, 10 g/L tryptone) from a frozen stock and incubating overnight at 37 °C, while shaking at 200 rpm. Approximately 30 μL of solution was then resuspended in 3 mL of the same medium and incubated under the same conditions for 3 h. To ensure a high percentage of motile bacteria in the experiments, non-motile and dead cells were gently removed from bacterial suspensions using sterile cell culture inserts incorporating a 3-μm-pore-size membrane, following a procedure described before ^12^ (Supplementary Information). To investigate the impact of the medium on the bacterial distribution, experiments were also carried out using TB diluted 1:10 in an isotonic saline solution (NaCl 5 g/L) and AB minimal medium. *P. aeruginosa* cells were centrifuged (1000*g* for 10 min) and then resuspended in the AB medium. The bacterial preparation procedures did not affect bacterial swimming speed or tumbling rate.

### Microfluidic assays

To analyze surface attachment in flow around a pillar, we fabricated a microfluidic device with four channels on the same chip, each containing six pillars: two each of diameter 50, 100 and 200 μm (Fig. 1a). In order to ensure that the dominant velocity gradients occurred near the pillars, we designed the microchannel with aspect ratio *H/W* = 0.1 (height *H* = 100 μm; width *W* = 1 mm). To analyze surface attachment in flow around pillars of different wettability, we fabricated a microfluidic device with four channels on the same chip. Two channels were plasma-treated and bonded 5 days before the experiment, and two were bonded just 1 hour before the experiment. Due to the PDMS hydrophobic recovery ^56^, surfaces with a very different contact angle were obtained (Fig. 6a; contact angle 95° ± 5° for the sample with 5 days of recovery time and of 15° ± 5° for that with 1 hour recovery, estimated from images of water droplets on the surface). To analyze surface attachment in flow around pillars of different stiffness, we fabricated a microfluidic device with six channels on the same chip. Stiffness was controlled by altering the concentration of the cross-linking agent in the PDMS solution (Fig. 6d). Pairs of channels were produced by adding a cross-linker concentration of 3%, 5% and 10% (typical concentration). All the channels were cured at 80 °C for 3 hours, bonded to glass, and then stored for 24 hours at ambient temperature to fully polymerize. According to literature ^57,58^, the PDMS has a Young modulus in the range of 150 ± 50 kPa with a concentration of 3%, of 500 ± 100 kPa with 5%, and of 2.25 ± 0.25 MPa with 10%. To analyze surface attachment in sinusoidal and corrugated channels, we fabricated two microfluidic channels with one flat wall and the other wall having either sinusoidal features (Fig. 7a) or irregularly corrugated features (Fig. 8). Potential confounding factors stemming from cell growth, cell–cell signaling, extracellular matrix production and biological variability were minimized by focusing on early attachment (<5 h) and by performing experiments at different shear rates in parallel using a single cell culture. Flow was driven by a syringe pump (neMESYS 290N, CETONI, Germany), using flow rates in the range *Q* = 0.6–6 μL/min. Prior to use, all the microfluidic channels were washed with 2 mL of medium. A diluted PA14 bacteria suspension (OD < 0.01; cell concentration < 10^6^ cells/mL) was flown for 5h. All experiments were performed at room temperature.

### Cell imaging and tracking

All imaging was performed on an inverted microscope (Ti-Eclipse, Nikon, Japan) using a digital camera (ORCA-Flash4.0 V3 Digital CMOS camera, Hamamatsu Photonics, Japan). Bacterial trajectories (Fig. 4a), were acquired using phase-contrast microscopy (30× magnification, 200 frames per second). Bacterial attachment (Fig. 1d) was quantified using epifluorescence microscopy (30× magnification, 6 images per hour). All image analysis was performed in Matlab (The Mathworks) using in-house cell tracking algorithms.

### Statistics and derivations

All images of bacterial attachment on pillars were taken at channel mid-depth after 5 h of continuous flow. The fluorescent image shown in Fig. 1d was acquired during a single experiment. The intensity distributions of the fluorescent signal shown in Fig. 2a,b,c were averaged over two identical pillars; the experiment was repeated three times with consistent results, but data shown are from a single realization. Data on experimental capture efficiency shown in Fig. 3a,c, on angular distributions of the fluorescence intensity shown in Fig. 5a-d, and on the normalized standard deviation *σ*_*θ*_/*σ*_*u*_ shown in Fig. 5e were obtained from experiments repeated at least three times, and error bars correspond to the standard error of the mean. Each value was averaged over at least six identical (*i*.*e*., same-diameter, same material) pillars. Sample trajectories of PA14 shown in Fig. 4a represent the longest trajectories obtained experimentally of bacteria colliding with the pillar; out of a total of 10^3^ trajectories recorded, 80 intercepted the pillar. Each of the angular distributions of the fluorescent intensity shown in Fig. 6b,e was averaged over four identical pillars. The surface coverage data shown in Fig. 6c,f were obtained as an average of the surface coverage measured in twelve images (450 μm × 450 μm) of the PDMS surface, at a location 2 mm upstream from each pillar and on the upper PDMS surface of the microchannel; error bars correspond to the standard error of the mean. The intensity distribution of the fluorescence intensity shown in Fig. 7b was averaged over 100 identical periods and normalized by the maximum intensity value. The fluorescent image shown in Fig. 8a was acquired during a single experiment.

### Quantification of the capture efficiency

In the model results, the capture efficiency *η*_C_^mod^ was calculated, according to its definition, as the fraction of bacteria removed from the volume of water subtended by the pillar’s cross-section ^28,32^. In the experiments, the capture efficiency *η*_C_^exp^ was obtained (see Eqn. (7) in ref. ^32^) by dividing the bacterial capture rate by the flux of particles encountering the cylinder, *F = PU d*_P_ *l*_P_, where *P* is the particle concentration in the flowing fluid and *l*_P_ is the height of the cylinder (here set to 1, since we are considering a 2D plan at channel mid-depth). Capture rate is assumed proportional to bacterial attachment, where the latter was measured in terms of fluorescent intensity^24^, *I*_IN_, of all pixels integrated over a 5-μm-thick annulus around the perimeter of the pillar. We thus define the experimental capture efficiency as *η*_C_^exp^ = *I*_IN_ / *(P U d*_P_ ).

### Numerical simulations

The mathematical model is based on first computing the 3D velocity field inside the microchannel (with COMSOL Multiphysics) and then using it to simulate the transport of individual bacteria, for the same geometry and flow conditions as in the experiments. We modeled bacteria as prolate ellipsoids with an effective aspect ratio *q*, which accounts for the combined hydrodynamic resistance of cell body and flagellar bundle, and swimming speed *V* directed along the cells’ long axis. A cell’s equations of motion in the 2D flow that occurs in the experimental observation plane (i.e., channel mid-depth; Fig. 1b) are

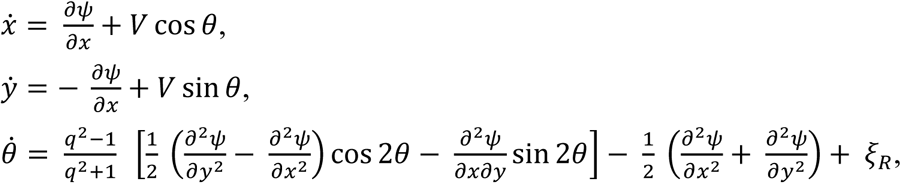

where *ψ* is the streamfunction and *ξ*_R_ is the rotational noise represented as a Gaussian-distributed angular velocity with mean zero and variance 2*D*_R_*/Δt*, where *Δt* is the elapsed time and *D*_*R*_ is the cell’s effective rotational diffusivity. The equations of motion were integrated numerically for up to 5 ×·10^5^ bacteria using a fourth-order Runge-Kutta scheme implemented in Matlab (The MathWorks). The swimming speed and rotational diffusivity were measured experimentally (Supplementary Fig. 1), while the effective aspect ratio, *q*, was computed from resistive force theory (Supplementary Information). For *P. aeruginosa* we used *q* = 9.4, *V* = 45 μm/s, and *D*_R_ = 1.4 rad^2^/s (Supplementary Fig. 2). For *E. coli* we used *q* = 8.5, *V* = 21.6 μm/s, and *D*_R_ = 0.32 rad^2^/s (Supplementary Information). For spherical non-motile bacteria (Fig. 6e,f) we used *q* = 1, *V* = *D*_R_ = 0. In the simulations, the swimmers start 400 μm upstream of the pillar (i.e., the beginning of the flow field) with random positions across the channel width (*y*) and random orientations (*θ*). Each time a swimmers contacts the pillar surface or exits the flow field it is re-injected with prescribed initial conditions. For contact between bacteria and surfaces, we used a ‘perfectly sticking’ condition, i.e., any collision between the simulated trajectory and the boundary of the pillar or of the corrugation is considered as the bacterium irreversibly attaching to the surface. This condition, which neglects any hydrodynamic or specific interaction between the cell and the surface of the pillar, represents a good approximation for the surface attachment of biofilm-forming bacteria, such *P. aeruginosa* and *E. coli*.

## Acknowledgements

The authors acknowledge support from an ETH Fellowship and an SNSF PRIMA grant (to E.S), from Gordon and Betty Moore Marine Microbial Initiative Investigator Award GBMF3783 (to R.S.), and from Simons Foundation Grant 542395 (to R.S.) as part of the Principles of Microbial Ecosystems Collaborative (PriME).

## Author contributions

E.S., R.R., G.L.M., V.K. and R.S. designed research. E.S. and R.R performed experiments on pillars, simulations for all geometries, and analyzed all the data. G.L.M. performed experiments on sinusoidal and wavy channels. A.V. and L.E. modified *Pseudomonas aeruginosa* strains. E.S., R.R. and R.S. wrote the manuscript and all authors edited and commented on the manuscript.

## Supplementary Information

### Culture preparation and media

The following procedure was used immediately prior to experiments in order to obtain motile cells, according to [1]. 1.5 mL of culture medium was added to a single well of a 6-well cell culture plate (BD Biosciences, San Jose, CA). A sterile cell culture insert (BD Biosciences, San Jose, CA), incorporating a polyethylene terephthalate (PET) track-etched membrane (pore size: 3 μm), was placed into the well and wetted with the culture medium, while avoiding trapped air under the membrane. 1 mL of cultured bacterial solution was then added to the insert. The bacterial culture was allowed to stand for 15 min, during which time motile bacteria migrated through the membrane, while non-motile cells settled onto the membrane surface. The pore size of the membrane was chosen to allow the passage of only single cells. The filter insert was gently removed and the remaining solution in the well, containing a high fraction of motile cells, was diluted to OD600 = 0.01 and used in experiments. AB medium was prepared according to the following protocol [2]: 200 mL of a 5xA solution (add 2 g of (NH_4_)_2_SO_4_, 6 g of Na_2_HOP_4_, 3 g of KH_2_PO_4_ and 3 g of NaCl to 200 mL of deionized water; mix and autoclave) and 800 mL of 1xB solution (add 1 mL of a 0.1 M CaCl_2_ sterile solution, 1 mL of a 1 M MgCl_2_ sterile solution and 1 mL of a 0.003 M FeCl_3_ sterile solution to 797 mL of sterilize deionized water).

### Bacterial GFP tagging

A green fluorescent protein (GFP) [3] was PCR amplified using pGFP_Fw_XbaI (tcctctagaGCGGCCGCTCTAGACATT) and pGFP_Rv_KpnI (ttcggtaccGTACCCAGCTGTTGACTCG). The amplicon was cleaned (QIAquick PCR Purification kit, Cat No./ID:28106), cut with KpnI_HF (New England Biolabs, Cat No. R3142S) and XbaI (New England Biolabs, Cat No. R0145S) and ligated into the plasmid pBBR1MCS-3 [4] digested with the same enzymes. To mark the *P. aeruginosa* PA14 wild-type and mutant strains (*motB* and *flgE*) [5] with GFP, plasmid pBBR1MCS-3::gfp was mobilized into each strain by triparental mating as described previously [6] using the helper *E. coli* pRK2013 [7].

### Microfluidic assay

Microfluidic channels were fabricated using standard soft lithography techniques. Microchannel molds were prepared by depositing SU-8 2150 (MicroChem Corp., Newton, MA) on silicon wafers via photolithography. Polydimethylsiloxane (PDMS; Sylgard 184 Silicone Elastomer Kit, Dow Corning, Midland, MI) was prepared (with 10% by weight of cross-linker) and cast on the molds. After curing, PDMS microchannels were plasma-sealed onto a clean glass slide. Channels were flushed with 2 ml of fresh medium before each experiment.

### Particles’ diffusion coefficient

The particles’ diffusion coefficient was computed as follows. For Brownian particles, the diffusion coefficient is given by the well-known Stokes-Einstein relation, i.e., *D* = *D*_*0*_ = *k*_*b*_*T/6πηR*, where *k*_*b*_ is the Boltzmann constant, *T* is the temperature of the system, *η* is the fluid viscosity and *R* is the particle radius. The random walk-like swimming behavior of motile bacteria allows the definition of an effective diffusion coefficient that is expressed as *D = D*_*0*_ + *(V*^*2*^ *τ*_*R*_*) / 4*, where *V* is the bacterial swimming speed and *τ*_*R*_ is the rotational diffusional time [8].

### Langevin model of bacterial motility in flow

We assumed that the fluid forces on bacteria were governed by the Stokes flow conditions. The effective aspect ratio of a *P. aeruginosa* cell, composed of a 2.4 μm by 1.2 μm ellipsoidal bacterial cell body and 10-μm-long helical flagellar bundle, used in the model was *q* = 9.4; the effective aspect ratio of a *E. coli* cell, composed of a 1 μm by 0.5 μm ellipsoidal bacterial cell body and 8-μm-long helical flagellar bundle, used in the model was *q* = 8.5. These values were computed from resistive force theory. Briefly, we computed the hydrodynamic resistance coefficients of the composite structure (cell and flagellar bundle) and then determined the aspect ratio, *q*, of an ellipsoid having the same rotational mobility in shear (i.e., the same Jeffery orbit). The cell swimming speed used in the model, *V* = 45 μm/s for *P. aeruginosa* (Supplementary Fig. 2) and *V* = 21.6 μm/s for *E. coli*, and rotational diffusivity, *D*_R_ = 1.4 rad^2^/s for *P. aeruginosa* and *D*_R_ = 0.32 rad^2^/s for *E. coli*, were directly measured by tracking individual cells in the absence of flow (for more details see [1]).

---

**Supplementary Figure 1.**
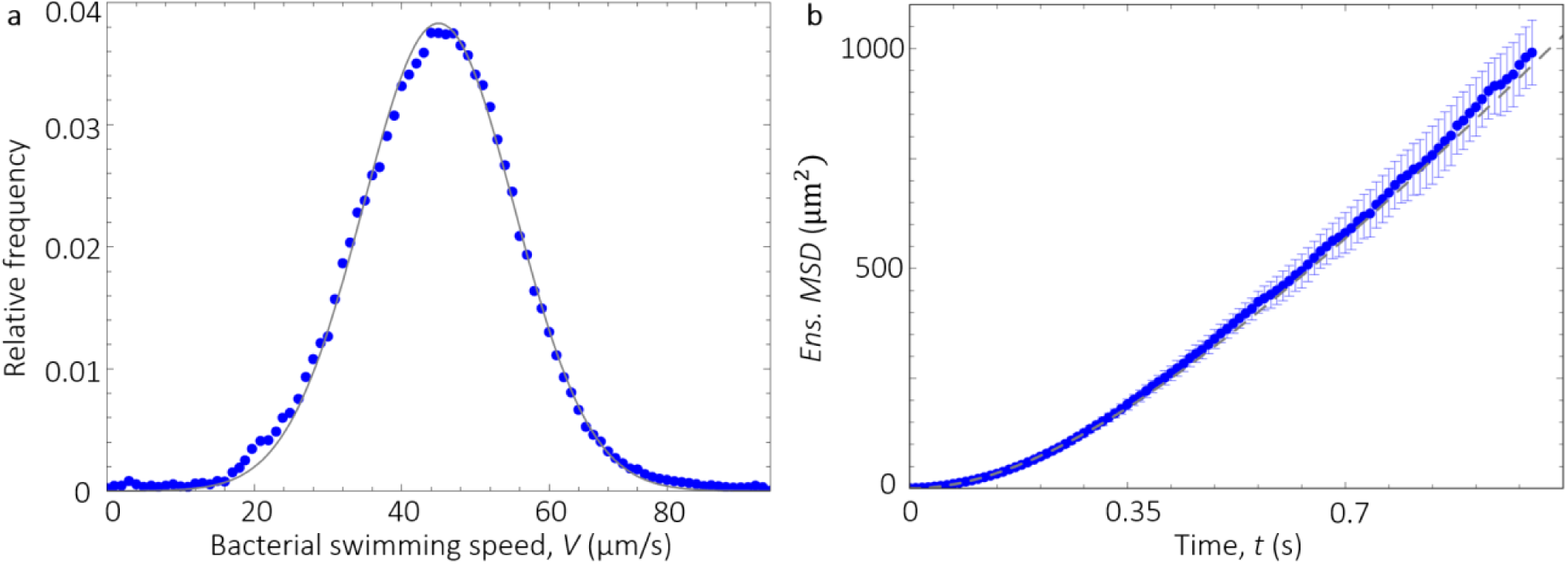
Measured swimming properties of *P. aeruginosa*. **a** Distribution of swimming velocities (blue circles) measured by tracking individual cells in a population of wild-type *P. aeruginosa*. The gray curve is a Gaussian fit, which yields a mean swimming speed of *V* = 45.02 μm/s with standard deviation 10.25 μm/s. **b** Ensemble mean-square displacement (*MSD*) for measured cells for the same bacterial population (blue circles). The ensemble average was computed over >1500 cells tracked during three different experiments and then averaged. The gray dashed curve is a fit using the formula *MSD* = 0.5 (*V* ^2^/*D*_R_^*2*^) [2*D*_R_*t* + *exp*(−2*D*_R_*t) -* 1], which yields a rotational diffusivity of *D*_R_ = 1.4 rad^2^ s^−1^.

**Supplementary Figure 2.**
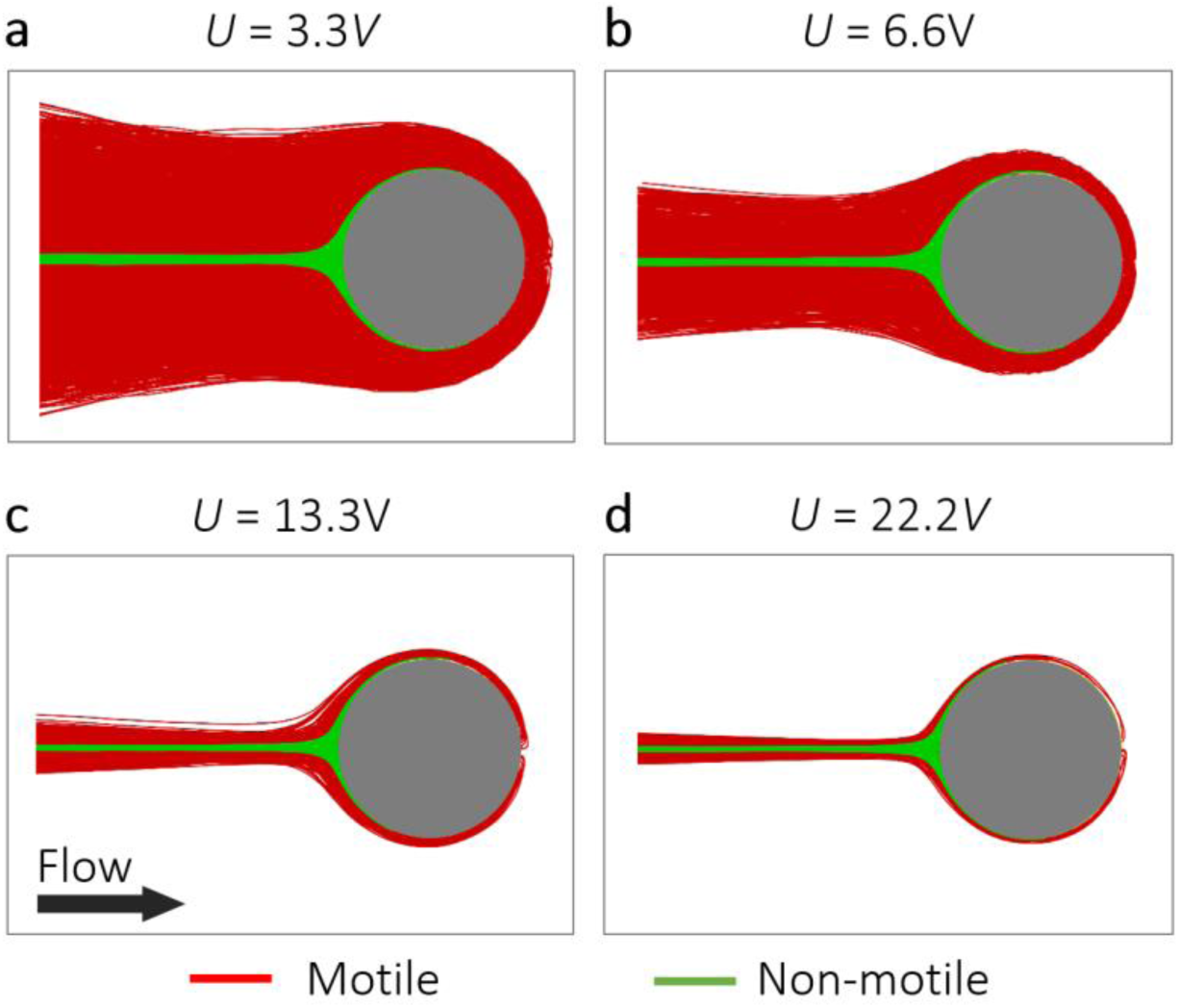
Bacterial motility dramatically increases the capture area in flow. **a-d** Trajectories that encountered the 100-μm pillar, obtained with the model for motile (red) and non-motile (green) cells for a mean flow velocity *U/V* = 3.3 (a), *U/V* = 6.6 (b), *U/V* = 13.3 (c) and *U/V* = 22.2 (d).

**Supplementary Figure 3.**
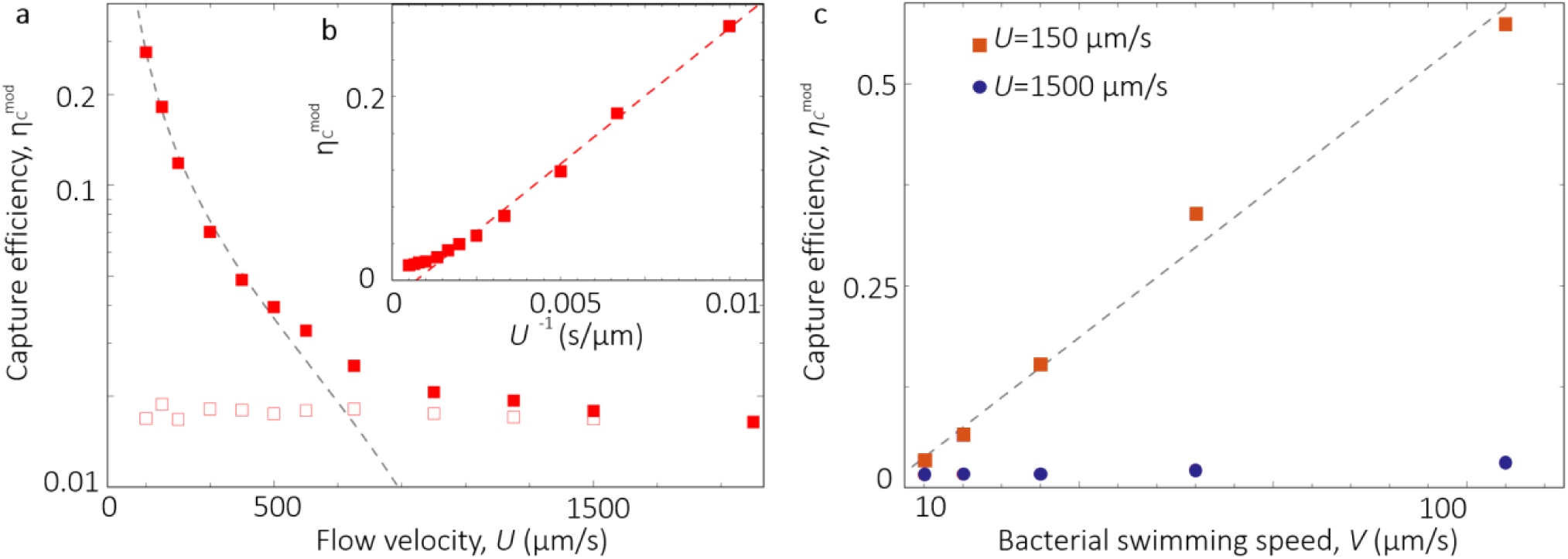
Capture efficiency as a function of flow velocity and bacterial swimming speed. **a** Capture efficiency, *η*_C_^mod^, as a function of the flow velocity, *U*, for a 100-μm pillar obtained from the model for motile (filled squares) and non-motile (open squares) cells. The gray dashed curve shows the scaling *η*_C_^mod^∼*U*^−1^. **b** Capture efficiency, *η*_C_^mod^, as a function of the inverse of the flow velocity, *U*^−1^. The red dashed curve shows the relationship *η*_C_^mod^∼*U*^−1^. **c** Capture efficiency, *η*_C_^mod^, as a function of the bacterial swimming speed, *V*, for a 100-μm pillar obtained from the model at two values of imposed mean flow velocity, *U*. The gray dashed curve shows the scaling *η*_C_^mod^∼ *V*.

**Supplementary Figure 4.**
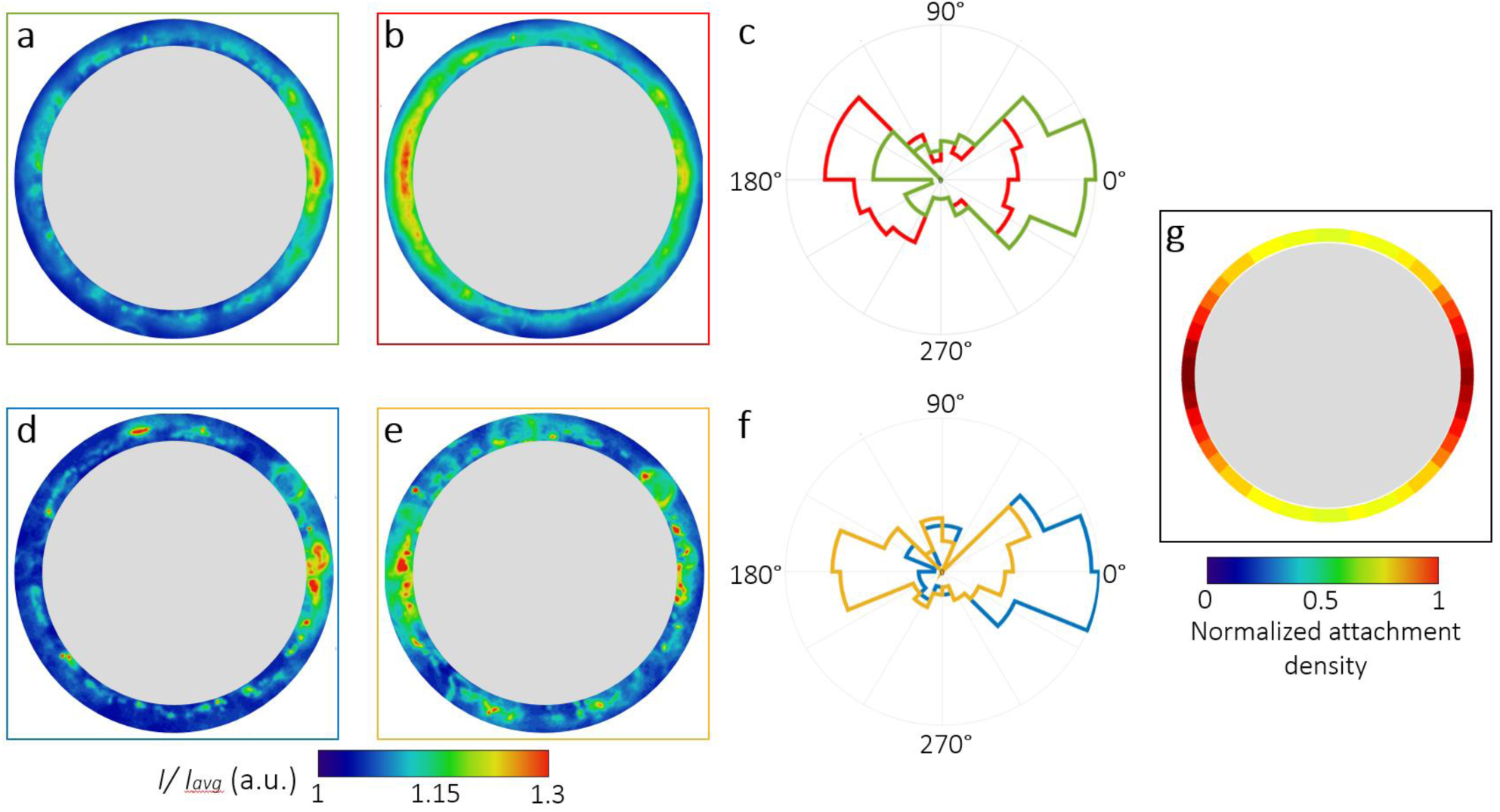
A flow reversal causes a reversal in the location of bacterial attachment. **a** Distribution of fluorescent intensity, *I*, normalized by its mean value, *I*_*avg*_, after 2.5 h of flow from left to right of a diluted suspension of PA14 wt GFP cells in Tryptone Broth, for *U/V* = 6.6 and a 100-μm-diameter pillar. Note the leeward attachment peak on the right side of the pillar. **b** Same, but after a further 2.5 h of flow from right to left. Note the leeward attachment peak on the left of the pillar. Each intensity distribution is the average over 12 identical pillars. **c** Angular distribution of the fluorescence intensity, *I*, for the cases shown in panels a (green) and b (red). **d, e** Same as a and b, respectively, but for cells in AB minimal medium. Each intensity distribution is the average over 6 identical pillars. **f** Angular distribution of the fluorescence intensity, I, for the cases shown in of the panels d (blue) and e (yellow). **g** Angular distribution of the normalized attachment density of bacteria on the pillar predicted by the mathematical model in the same flow conditions as panel b, *i*.*e*., 2.5 h of left-to-right flow followed by 2.5 h of right-to-left flow. Simulations confirm that flow reversal causes a reversal in the location of bacterial attachment.

**Supplementary Figure 5.**
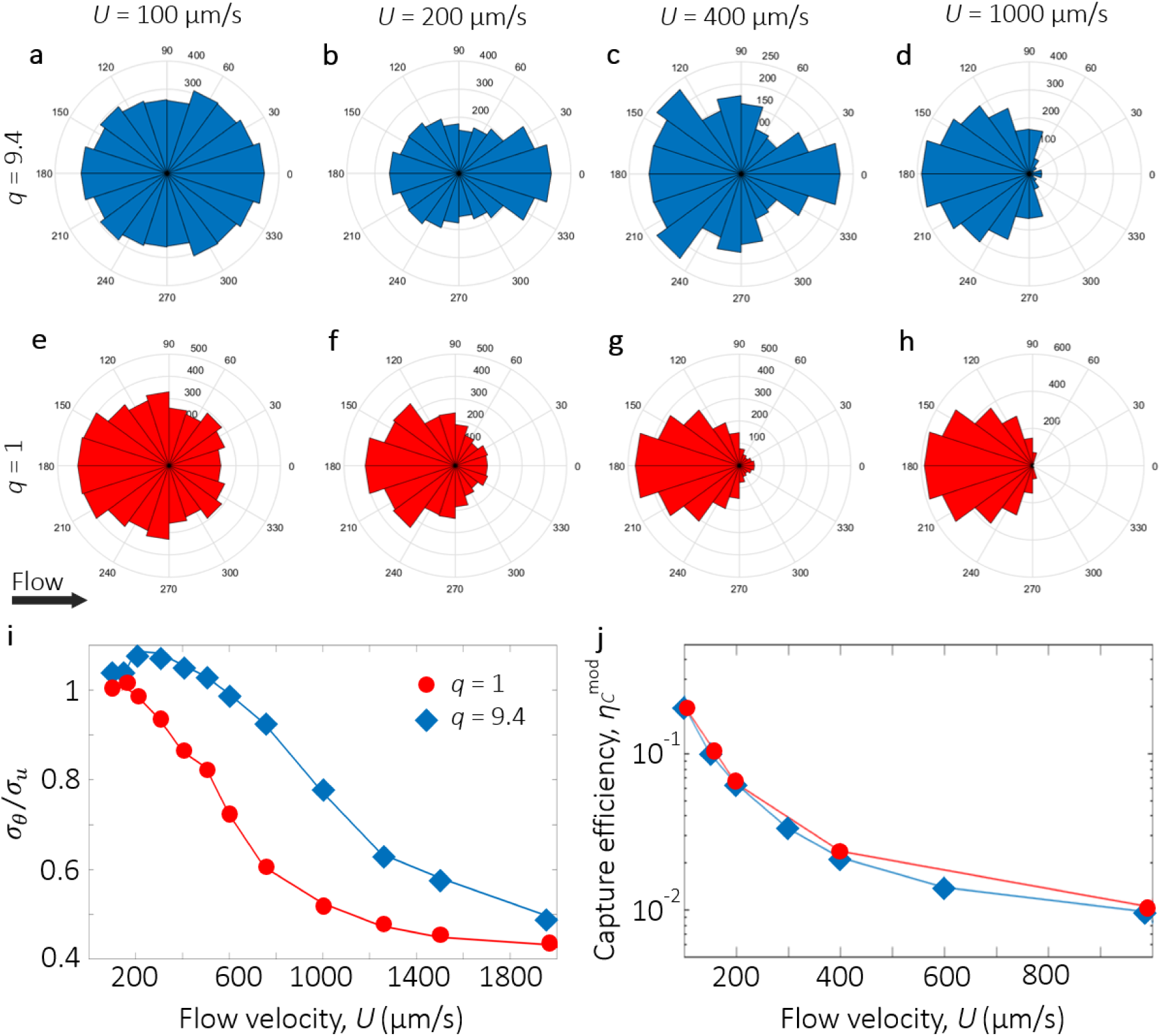
Attachment distribution and capture efficiency for elongated and spherical motile bacteria. **a-h** Polar distribution of the attachment density of motile elongated (*q* = 9.4; a-d) and spherical (*q* = 1; e-h) cells on a 100-μm pillar at mean flow velocity of 100 μm/s (2.2*V;* a, e), 200 μm/s (4.4*V;* b, f), 400 μm/s (8.8*V;* c, g) and 1000 μm/s (22.2*V;* d, h). **i** Normalized standard deviation *σ*_*θ*_/*σ*_*u*_ obtained with the model as a function of the mean flow velocity, *U*, for the100-μm pillar for motile elongated (*q* = 9.4; blue diamonds) and for motile spherical cells (*q* = 1; red circles). **j** Capture efficiency, *η*_C_^mod^, as a function of the mean flow velocity, *U*, for a 100-μm pillar obtained from the model for motile elongated (*q* = 9.4; blue diamonds) and for motile spherical cells (*q* = 1; red circles).

**Supplementary Figure 6.**
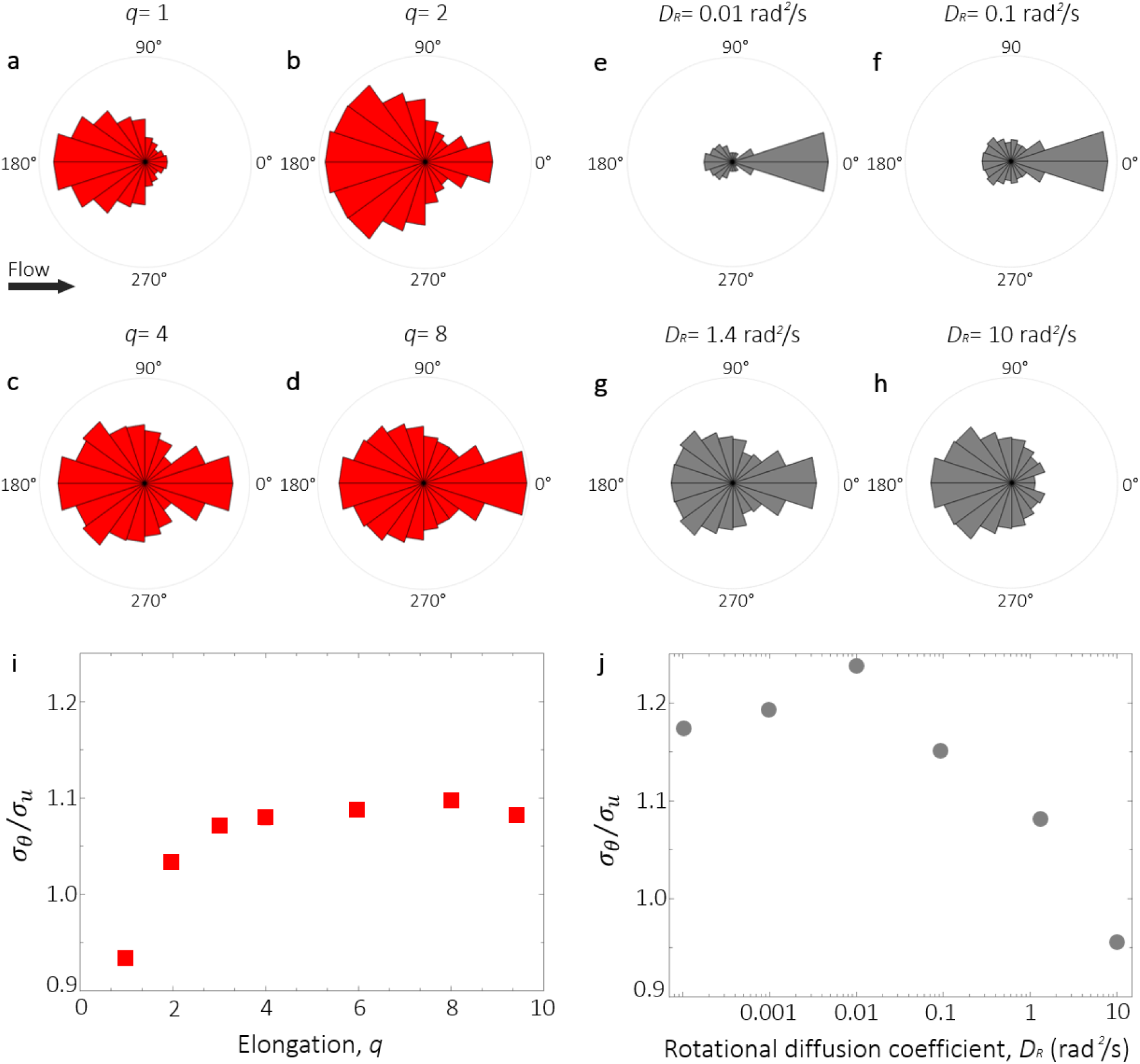
Angular distribution of the attachment of motile bacteria with different elongations and rotational diffusivities, predicted by the mathematical model for a relative flow speed of *U/V* = 6.6 and a 100-μm-diameter pillar. **a-d** Polar distribution of the attachment density of cells with elongation *q* = 1 (a), 2 (b), 4 (c) and 8 (d). **e-h** Polar distribution of the attachment density of cells with rotational diffusivities *D*_*R*_ = 0.01 rad^2^/s (e), 0.1 rad^2^/s (f), 1.4 rad^2^/s (g) and 10 rad^2^/s (h). **i, j** Normalized standard deviation of the polar distribution, *σ*_*θ*_/*σ*_*u*_, as a function of (i) the cell elongation *q* and (j) the cell rotational diffusivity *D*_*R*_.

**Supplementary Figure 7.**
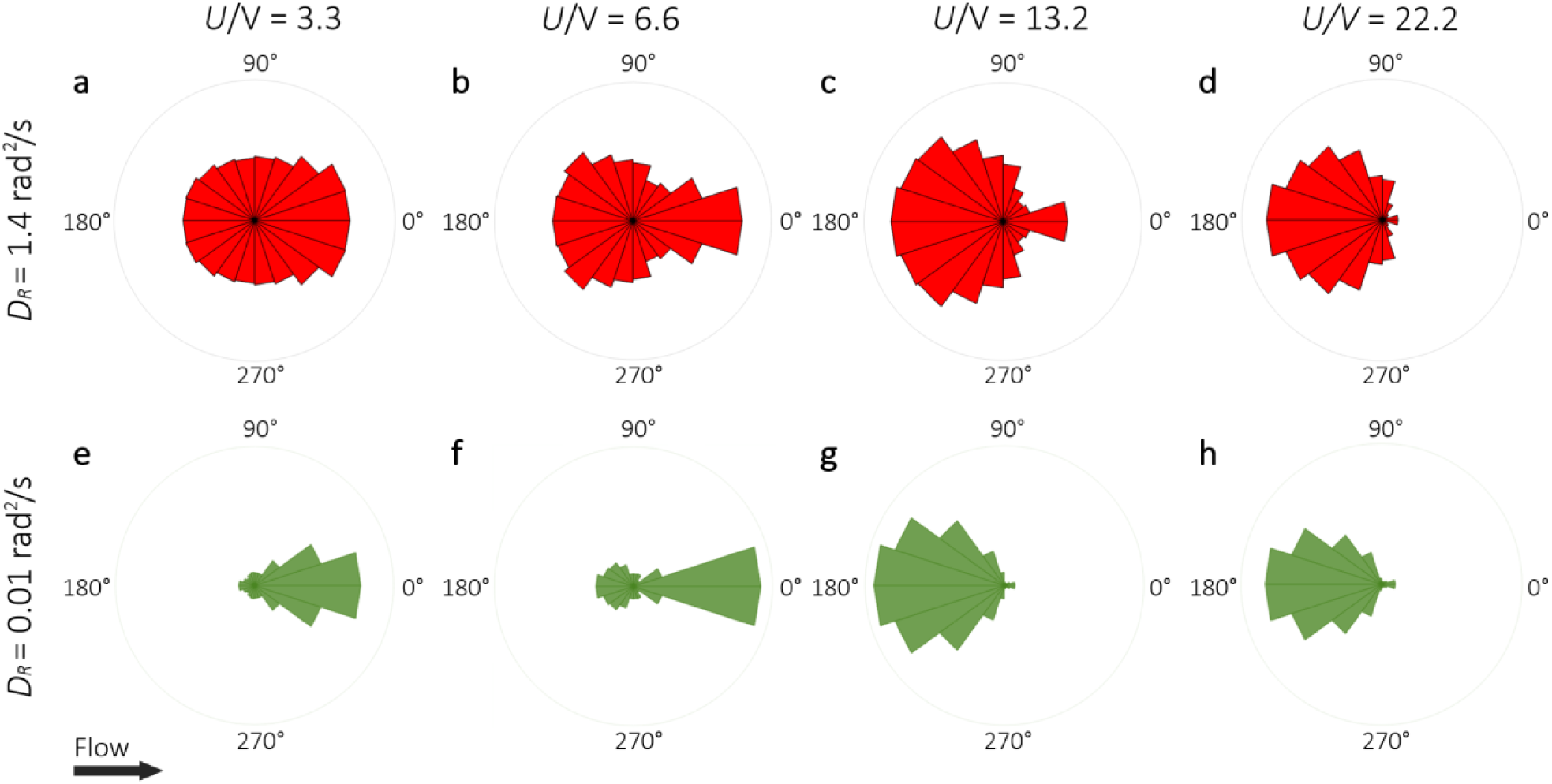
Angular distribution of the attachment of motile bacteria for different flow speeds *U*/*V*, and two values of the rotational diffusivity *D*_*R*_ predicted by the mathematical model for a 100-μm-diameter pillar and for cells with elongation *q* = 8.

**Supplementary Figure 8.**
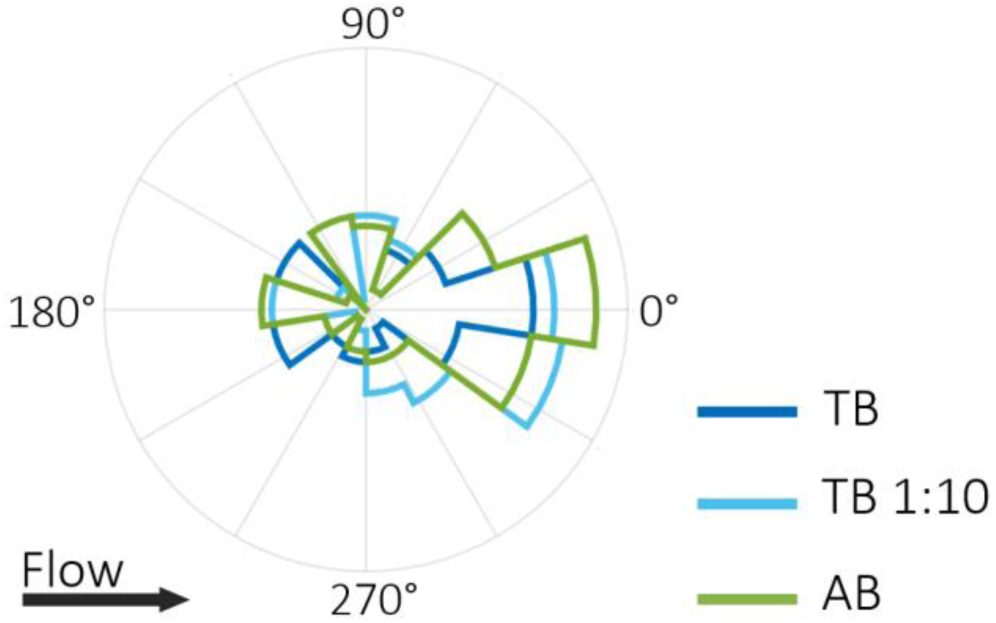
The culture medium does not affect the angular distribution of bacterial colonization around a pillar. Angular distribution of the fluorescence intensity, *I*, on a 100-μm pillar after 5 h of flow at *U/V* = 6.6 of a diluted suspension of PA14 wt GFP cells in different culture media: Tryptone Broth (TB; blue), Tryptone Broth diluted 1:10 in an isotonic saline solution (TB 1:10, cyan) and AB medium (AB, green). Each intensity distribution is the average over 12 identical pillars.

**Supplementary Figure 9.**
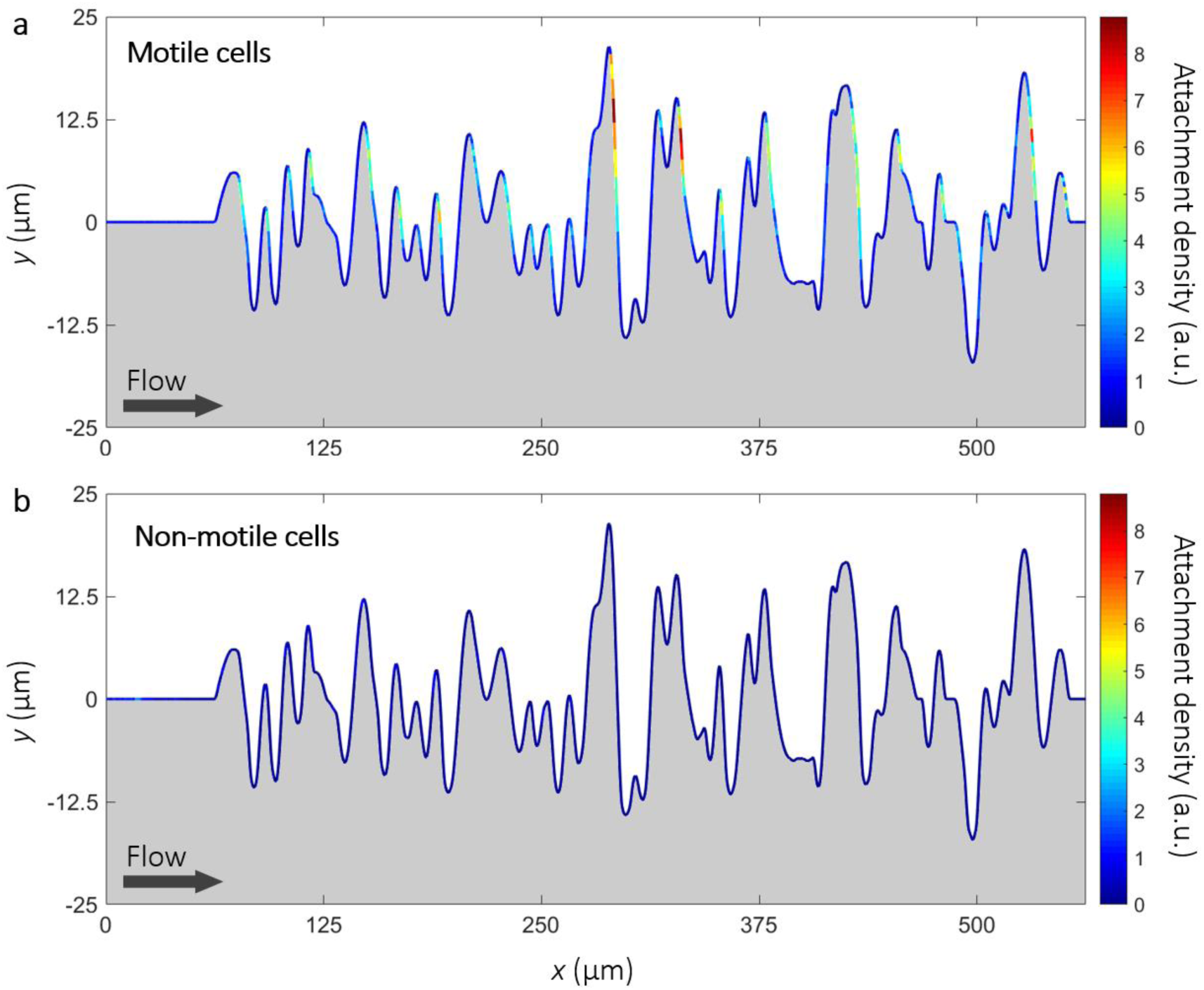
Colonization of a random corrugated surface by motile and non-motile cells. Normalized attachment density of motile (*q* = 8.5, *V* = 21.6 μm/s; a) and non-motile (*q* = 8.5, *V* = 0 μm/s; b) cells on a random corrugated surface obtained with the model at a mean flow velocity of 150 μm/s.

